# Age related defects in NK cell immunity revealed by deep immune profiling of pediatric cancer patients

**DOI:** 10.1101/2020.03.09.983288

**Authors:** Eleni Syrimi, Naeem Khan, Paul Murray, Carrie Willcox, Tracey Haigh, Benjamin Willcox, Navta Masand, Jianmin Zuo, Sierra M Barone, Jonathan M Irish, Pamela Kearns, Graham S Taylor

## Abstract

Systemic immunity plays an important role in cancer immune surveillance and therapy but there is little detailed knowledge about the immune status of healthy children or children with cancer. We performed a high dimensional single cell analysis of systemic immunity in pediatric cancer patients and age-matched healthy children. In young children with cancer (age < 8years) NK cells were decreased in frequency, maturity, expression of perforin and granzyme-B, and were less cytotoxic in ex vivo assays. NK cell activity was restored after in vitro culture with interleukin-2. In contrast, older children with cancer (>8 years old) had decreased naive CD4 and CD8 T-cells with concomitant increases in effector memory and T effector memory RA-revertant (TEMRA) T-cells. These immunological changes in pediatric cancer patients are relevant to the better understanding of how cancers diagnosed in childhood interact with systemic immunity and could inform the development and application of effective immune-modulating therapies in the pediatric population.

**One Sentence Summary:** High dimensional analysis of systemic immunity in pediatric cancer patients reveals clinically relevant immune changes in NK and T-cells that vary with patient age.

## Introduction

Pediatric cancers are a major cause of childhood mortality ^1^. Overall, the incidence of these cancers shows a bimodal age-specific pattern with peaks between 1-5 years and 10-14 years of age ^1^. Current treatments are heavily reliant on chemotherapy and as such are associated with significant clinical challenges due to acute toxicity and often fail to achieve cure for patients with high risk, metastatic or relapsed disease ^2^. In addition, survivors suffer multiple long-term treatment-related sequelae and excess mortality due to cardiotoxicity, secondary cancers and infections ^3^. Improving cure rates, while decreasing side effects, requires new treatment strategies and harnessing the immune system is one possibility. Several immunotherapies, including CTLA4 and PD1 checkpoint inhibitors, are now licensed for some adult cancers, however for pediatric cancers emerging evidence suggests response rates for checkpoint inhibitors are low ^4^.

Systemic immune responses are essential for effective immunotherapy ^5^ and age-related immune variations ^6^ could contribute to differences in immunotherapy responsiveness. Current understanding of the immune systems of healthy children is limited. Most work has focused on neonates due to the ready availability of cord blood ^6^. The T-cell compartment of neonates is immature, tolerogenic and anti-inflammatory with reduced pre-inflammatory T helper (Th)1 function and skewing towards Th2 responses ^7^. Natural Killer (NK) cells are also present, but their cytotoxicity is less than half that of adult NK cells ^8^. A recent longitudinal analysis of immunity in neonates and children up to three months of age shows substantial changes occur in multiple immune cell types that follow a common trajectory ^9^. At three months of age the infant immune system has not reached the adult state. Additional immune development must therefore continue to occur throughout childhood, but this process has barely been characterized.

Cancer associated changes in systemic immunity are well documented in adults. Some, such as increases in the frequency of immunosuppressive cells like CD4+ regulatory T-cells (T-regs), are associated with a worse prognosis ^10^. Other changes, such as altered T-cell differentiation status, are associated with increased clinical response after immunotherapy ^11^. NK cells in adult cancer patients are unaltered in frequency but are less cytotoxic ^12^. This has been ascribed to soluble NK ligands released by cancer cells binding to NK receptors, thus inhibiting recognition of target cells by NK cells ^13^. It is not known whether these changes occur in children with cancer.

Differences also exist in the biology of cancers that afflict children and adults. Pediatric cancers have relatively few somatic coding mutations ^14^ meaning most patients’ tumors lack actionable neoantigens. Furthermore, cancers arising in young children (<5 years old) are often embryonal in origin. A cardinal feature of such tumors, which include hepatoblastoma ^15^, medulloblastoma^16^, neuroblastoma ^17^ and Wilms tumor^18^, is low or no expression of MHC-class I, making them poor T-cell targets. However, this loss could present a different therapeutic opportunity by rendering tumors sensitive to NK cell mediated lysis. In vitro experiments using NK cells derived from healthy adults show that neuroblastoma ^19^ and hepatoblastoma ^15^ cell lines and primary neuroblastoma tumor cells are all sensitive to NK-mediated lysis. Mass cytometry has advanced immunology research by allowing more than 40 markers to be simultaneously measured on millions of individual cells, allowing deep immunophenotyping to be performed with small volumes of blood ^20^. Partnered with innovative computational algorithms, such as tSNE ^21^, uMAP ^22^, flowSOM ^23^ and Marker Enrichment Modelling ^24^, mass cytometry has provided new insights into human immunology. In this study, we use mass cytometry to perform the first high dimensional single cell analysis of systemic immunity across several childhood cancers, comparing patients to age-matched healthy children. We identified important changes in the frequency and phenotype of multiple immune cell types, including T-cells and NK cells, and show many of these changes are influenced by patient age. These data provide novel insights into pediatric immunity in the context of malignancy and a rational basis on which the development of immunotherapies for these patients can be optimized.

## Results

### Frequency of NK cells and monocytes are altered in pediatric cancer patients

Pre-treatment peripheral blood mononuclear cells (PBMCs) from 20 children diagnosed with a range of childhood cancers and 19 age-matched healthy children were analysed using a 38-marker mass cytometry antibody panel (Supplementary Table 1). Demographic and clinical information are provided in Supplementary Table 2 and Supplementary Figure 1. One patient, C19, had pre-malignant nephroblastomatosis due to the inherited cancer predisposition Wilms tumor aniridia syndrome (WAGR) ^25^.

To aid visualisation of high-dimensional data, flow cytometry standard (fcs) files from patients and controls were concatenated into two separate files. Equal numbers of cells per individual (4751 cells) were used. After manually gating cells into two populations (i. CD3-CD19-, ii. CD3+ or CD19+ cells; Figure 1A) data were visualized using UMAP dimensionality reduction ^22^. The UMAP plots comparing patients and control donors (Figure 1B) showed clear differences in the frequency of innate immune cells, with monocytes increased and NK cells decreased in patients. Also evident were clear differences in the spatial distribution of immune cell populations on the two dimensions of the plot, particularly for CD4+ T-cells and NK cells, indicating that phenotypic variation existed between patients and controls. Analysing cell frequency per individual (Figure 1C) revealed patients had a significantly lower frequency of NK cells (p=0.0035) and a strong trend for increased monocytes (p=0.032). The frequency of T and B cell subsets were not significantly different.

**Figure 1.**
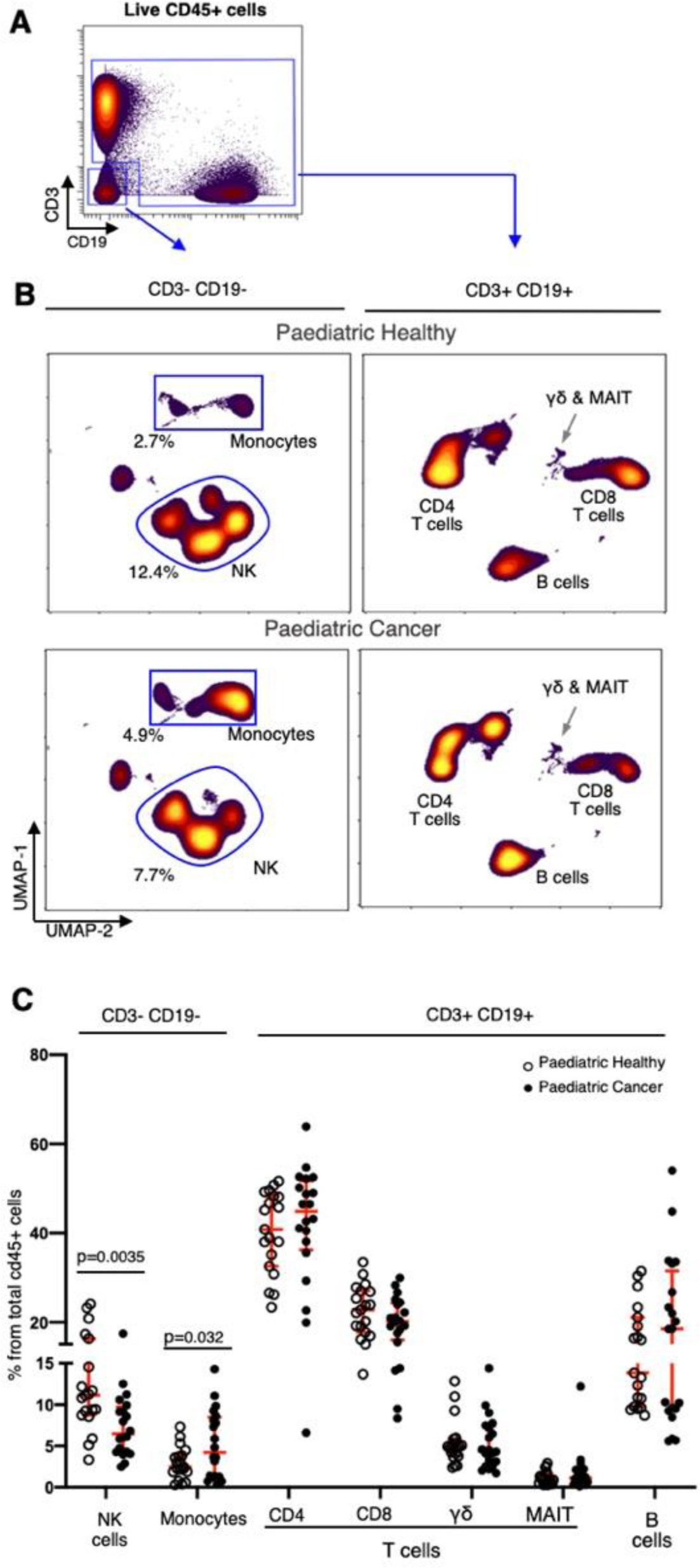
NK and monocyte cell frequencies are altered in pediatric cancer patients. A) Manual gating strategy used to split immune cells into CD3-CD19- and CD3+Cd19+subsets. (B) Concatenated fcs files for both pediatric healthy (n=19) and pediatric cancer patients (n=20) were analysed by UMAP dimensionality reduction. The frequency and identify of main immune subsets is shown. C) Frequency of the main immune subsets for all healthy children (open symbols) and cancer patients (closed symbols) expressed as a percentage of total CD45+ cells. The mean +/- 1 standard deviation is shown in red. P values were calculated using an unpaired t test, correcting for multiple comparisons (5% FDR).

### Classical monocytes are enriched in pediatric cancer patients

Given the frequency changes observed between patients and controls (figure 1), Marker Enrichment Modelling (MEM) ^24^, an algorithm that quantifies the enriched features of a cell population was applied in order to determine whether the monocytes were also altered in phenotype (Gating strategy shown in SI figure 2). This analysis showed monocytes from controls were positively enriched for HLA-DR (▴HLA-DR^+6^) whereas monocytes from patients were negatively enriched (▾HLA-DR^-4^, Figure 2A, left panel). Differences in monocyte HLA-DR levels were confirmed by measuring the median mass intensity on monocytes gated from each individual (Figure 2A, right panel, p=0.057).

**Figure 2.**
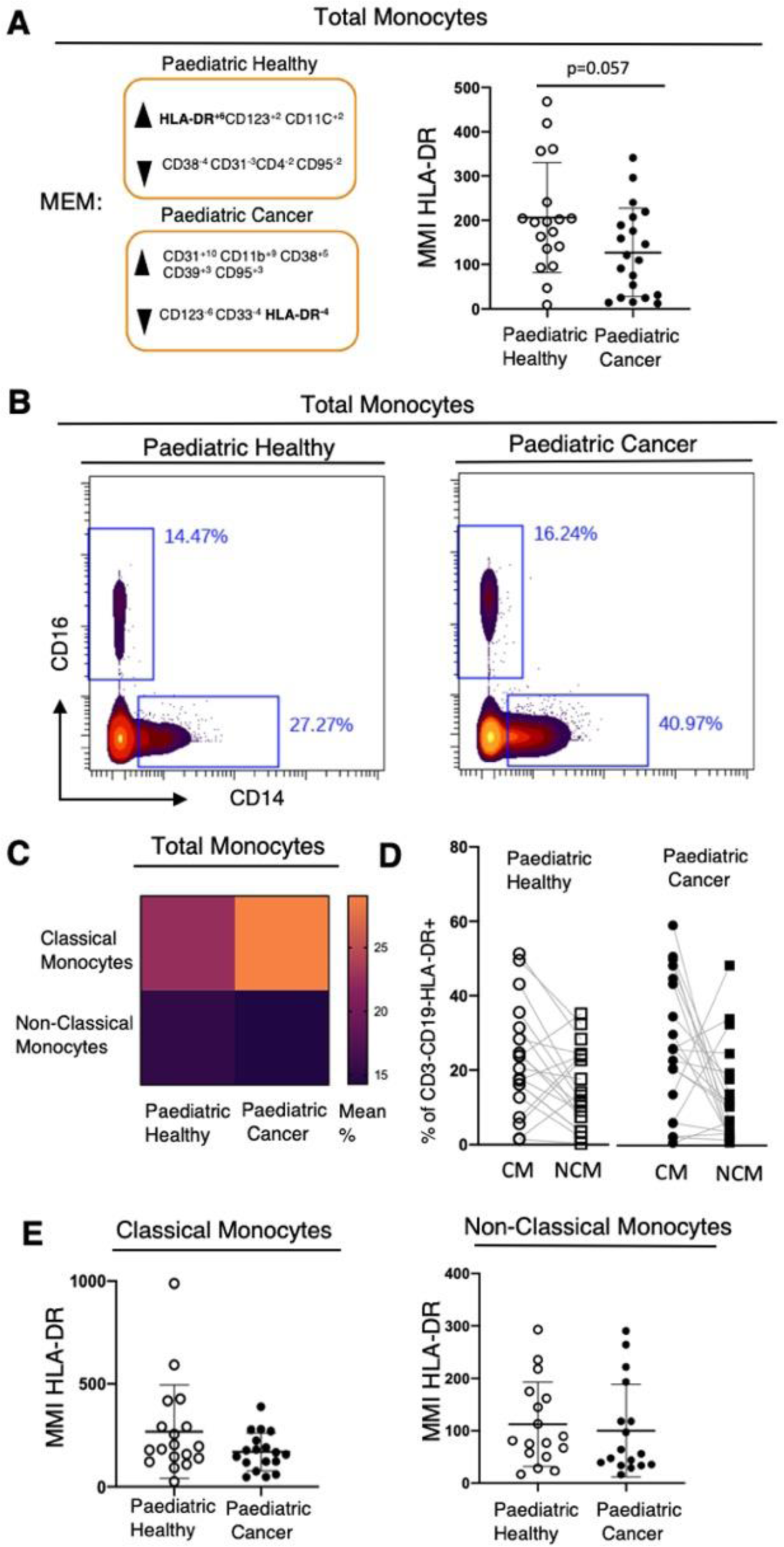
Characterisation of monocytes in pediatric cancer patients and controls. A) Left panel - results of marker enrichment modelling (MEM) of total monocytes. HLA-DR was negatively enriched in pediatric cancer patients. Right panel - median mass intensity of HLA-DR on monocytes gated for each individual showed decreased HLA-DR expression for patients (p=0.057, Mann-Whitney test). B) Manual gating strategy using CD14 and CD16 to identify classical monocytes (CM) and non-classical monocytes (NCM) in concatenated FCS files from pediatric healthy and cancer donors. C) Heatmap of mean abundance of CM and NCM in patients and healthy control donors. D) Frequency of CM and NCM for individual healthy control donors (open symbols) and cancer patients (closed symbols) expressed as a percentage of total monocytes. E). Median mass intensity of HLA-DR on CM (left panel) and NCM (right panel). Error bars represent mean +/- 1 standard deviation.

Classical monocytes (CM) and non-classical monocytes (NCM) differ functionally, being pro- and anti-inflammatory respectively, and also differ in HLA-DR expression ^26^. Using CD14 and CD16 to delineate each subset —CM (CD14+ CD16-) and NCM (CD14-CD16+) —we found patients’ monocytes contained a larger proportion of CM (Figure 2B and 2C). The CM:NCM ratio was 8.5 for patients compared to 2.4 for controls (Figure 2D). Four patients (C8, C9, C12, C16) had high CM:NCM ratios ranging from 12 to 54 – all four were younger than two years of age. Next, comparing HLA-DR levels on each monocyte subset, we observed HLA-DR levels on NCM were similar for patients and controls whereas there was a trend for patients’ CM to express lower surface levels of HL-DR (Figure 2E).

### NK cells in patients are shifted to an immature phenotype

Having observed differences in NK cell frequency and distribution on UMAP plots (Figure 1) we examined the NK cell phenotype in more detail. The classical NK differentiation markers CD56 and CD16 were used to divide NK cells into the four standard subsets ^27^. In order of increasing maturity these comprise subset 1 (CD56^bright^ CD16^-^), subset 2 (CD56^dim^ CD16^-^), subset 3 (CD56^+^ CD16^+^) and subset 4 (CD56^-^ CD16^+^). Analysis of the concatenated fcs files showed the frequency of immature subset 1 NK cells was 1.4-fold higher in patients compared to age matched controls (Figure 3A; 11.4% patients,4.8% controls,). This increase was apparent when heat plots of cell density from subset 1 were overlaid on black and white contour t-SNE plots of concatenated files gated on total NK cells (Figure 3B). Analysing these data at the level of individuals (Figure 3C) showed the increase in subset 1 immature NK cells in patients was statistically significant (p=0.0004). In contrast, the more mature NK subsets 2 and 4 were similarly abundant in patients and control donors and subset 3 (CD56^+^ CD16^+^) was slightly decreased in patients (0.1 fold decrease).

**Figure 3.**
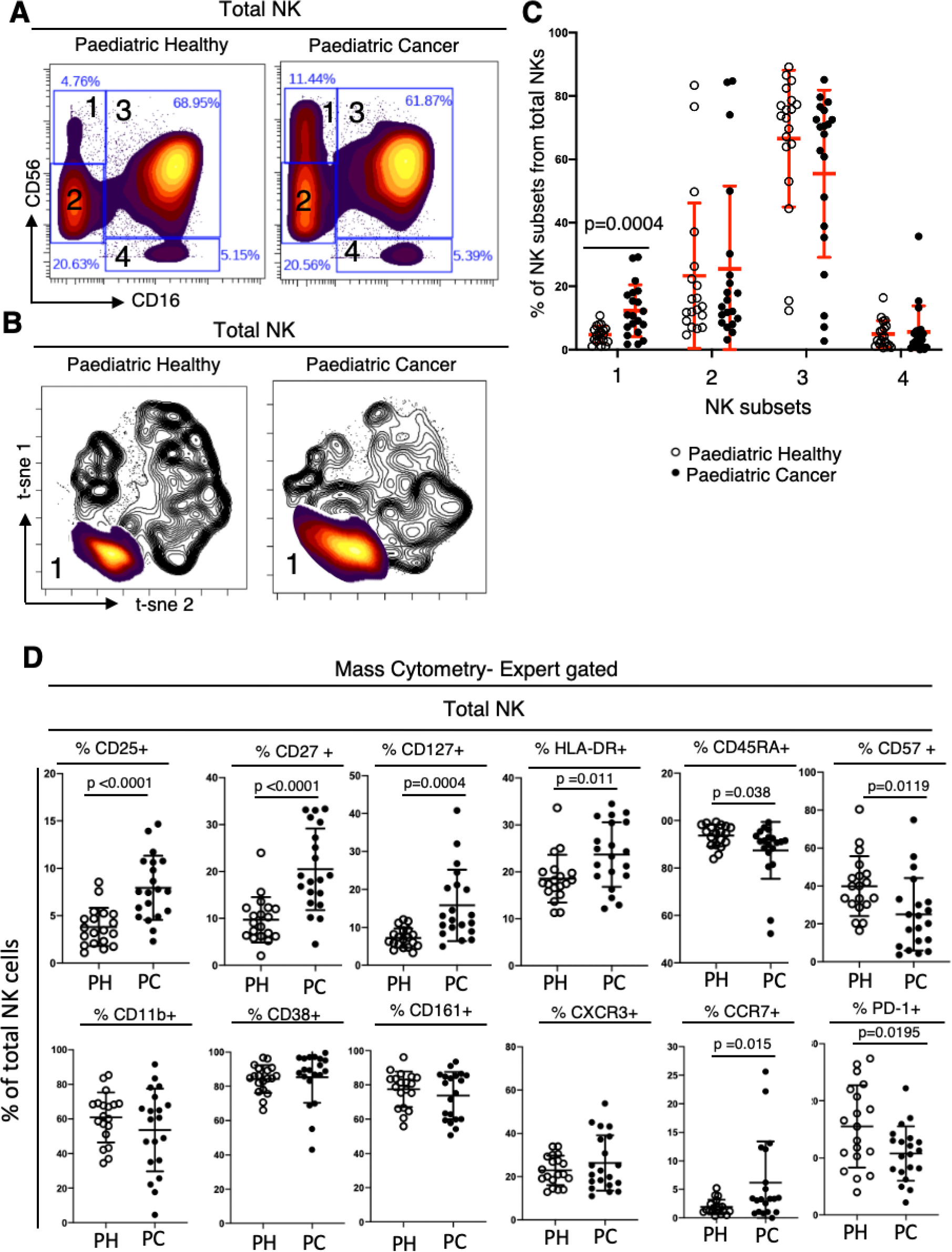
Detailed characterisation of NK cells in pediatric patients and controls. A) Biaxial plots of concatenated FCS files from healthy children (n=19) and cancer patients (n=20). Total NK cells were manually gated using CD16 and CD56 expression to delineate NK cells into four canonical differentiation subsets. In order of maturity: subset 1 (CD56^bright^CD16^-^), 2 (CD56^dim^CD16^-^), 3 (CD56^+^CD16^-^) and 4 (CD56^-^CD16^+^). The proportion of NK cells in each subset is shown on each plot. B) Heat plots of cell density from subset 1 were overlaid on black and white contour tSNE plots of total NK cells from pediatric healthy donors (left panel) and cancer patients (right panel). C) Frequency of each NK subset, expressed as a percentage of total NK cells per individual. The mean +/- 1 standard deviation is shown in red. There was no significant difference in frequency of subset 2, 3, or 4; subset 1 was significantly different (p=0.0004, unpaired t-test adjusted for multiple comparisons with 5% FDR). D) Mass cytometry analysis of pediatric cancer patients (PC,n=20) and age matched pediatric healthy (PH,n=19) showing percentage of total NK cells positive for the indicated markers.

Examining a broader range of NK phenotypic markers (Figure 3D) confirmed these differences in maturity. Patients had a significantly higher proportion of NK cells bearing markers associated with NK immaturity (CD25, CD27, CD127 and HLA-DR) ^28,29,30,31^ and a significantly lower proportion of NK cells bearing markers associated with NK maturity (CD45RA and CD57)^32,33^. No difference was observed in the proportion of NK cells expressing the adhesion molecule CD11b ^34^, the adhesion molecule and ectoenzyme CD38 ^35^ nor CD161, expressed by a distinct subtype of pro-inflammatory NK cells ^36^. There was also no difference in expression of the chemokine receptor CXCR3 which is required for NK cell accumulation in tumors ^37^. Finally, we examined the expression of the immune checkpoint PD1, which is expressed on a minority of mature NK cells in healthy adults and increased in adults with cancer ^38^. In contrast, our data showed that children with cancer had a significantly lower frequency (p=0.0195) of PD1-positive NK cells compared to their healthy age-matched counterparts. Examining PD1 expression on the four different NK cell subsets showed this difference was not due to patients having more immature NK cells (SI figure 3).

To further explore the NK cell phenotype we used a separate NK-focused mass cytometry panel (SI Table 3) to compare total NK cells from six patients and six age-matched controls (all selected from the originally analysed cohort). The inhibitory receptor NKG2A ^39^ was expressed on a significantly higher proportion of NK cells from patients (p=0.0325). There was no difference in the frequency of cells positive for the activatory receptors NKG2D, NKp46, NKp30 or DNAM1 ^39^ (Figure 4A). Examining each NK subset in turn, we found more frequent expression of NKG2A on mature cytotoxic NK cells (subset 3, SI fig 4; p=0.0018).

**Figure 4.**
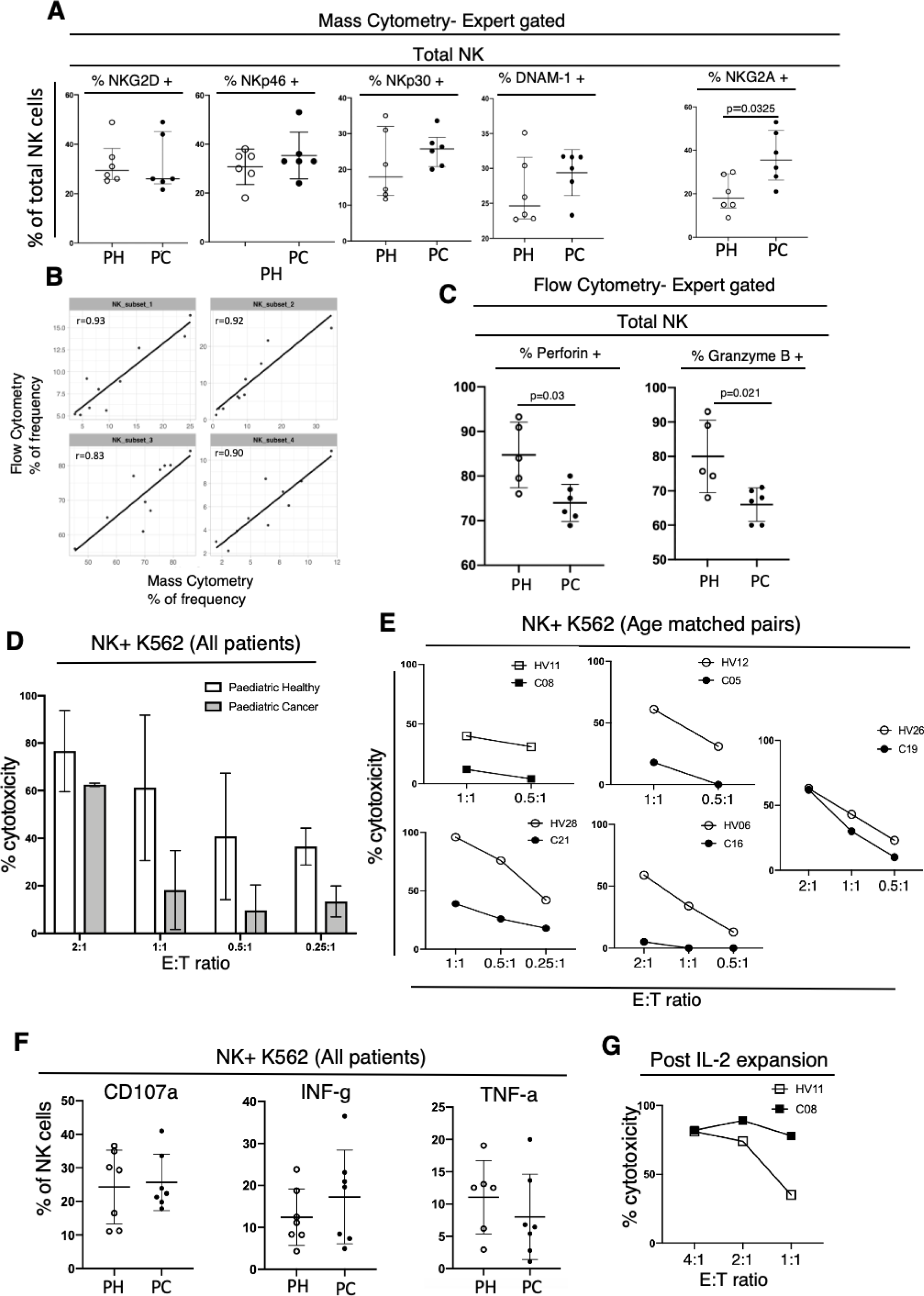
NK cells from cancer patients show decreased lysis but not decreased recognition of K562 target cells. Mass cytometry analysis of a subset of the patients and controls shown in Figure 3, now investigating NK receptors. Pediatric healthy (PH, n=6); pediatric cancer (PC, n=6). B) Comparison of NK subsets using mass and fluorescent cytometry demonstrated concordance. Pearson correlation coefficient for each subset is shown. C) Intracellular fluorescent flow cytometry showing percentage of total NK cells positive for perforin or granzyme B. D) Combined results of K562 cytotoxicity assays for five patients and five age matched control donors performed at the indicated NK effector to K562 target ratio. Mean cytotoxicity +/- 1 standard deviation is shown. E) Results from panel D now showing each patient and their age matched control. F) Flow cytometry analysis of NK cells from seven pediatric healthy donors (PH) and seven pediatric cancer patients (PC), which include patients analysed in D and E, measuring degranulation by CD107a as well cytokine production. Error bars shown mean +/- 1 standard deviation. F) K562 cytotoxicity assay performed using NK cells from patient C08 and their age matched control HV11 using NK cells expanded in vitro with IL-2 for 14 days. E:T indicates the effector: target ration used. For panels A-C results of unpaired t-tests, correcting for multiple comparisons with 5% FDR when necessary, are shown on the figure.

### NK cells from pediatric cancer patients are less cytotoxic in functional assays

Tumor surveillance and clearance by NK cells is mainly dependent upon their cytotoxic functions ^40^. We used fluorescent flow cytometry to measure levels of the intracellular cytotoxic effector molecules perforin and Granzyme-B (SI Table 4). Control experiments showed NK subsets identified by fluorescent cytometry were concordant with mass cytometry (Figure 4B). Testing samples from six patients and five age matched controls (all selected from the originally analysed cohort) we found that the total NK cell population in patients contained significantly lower proportions of cells positive for perforin (p=0.03) and cells positive for granzyme-B (p=0.02) (Figure 4C). This was not attributed to any particular NK cell subset (SI Fig5). We next measured the cytotoxicity of total NK cells using as a target the standard K562 cancer cell line ^41^. In these assays the effector to target ratio was adjusted to correct for the lower frequency of NK cells in patients. Results from five patients and five age-matched healthy controls showed that, overall, NK cells from patients were less cytotoxic (Figure 4D). NK cells from one patient, C19, showed similar levels of cytotoxicity to their age-matched counterpart (Figure 4E). Interestingly this patient had pre-malignant nephroblastomatosis rather than an established malignancy. To determine whether decreased K562 killing by patients’ NK cells was due to inefficient recognition or cytotoxicity, the assays were repeated using alternative readouts of NK cell function. Following co-culture with K562 cells, NK cells from patients and controls showed equivalent levels of degranulation and cytokine production (Figure 4F) showing recognition of NK cells was unaltered. In adult cancer patients soluble MICA and ULBP2 are elevated in the plasma and inhibit NK function ^42^. Unexpectedly, our pediatric cancer patients had lower plasma concentrations of MICA and ULBP2 compared to their healthy age matched controls (SI Fig 6). Collectively our results demonstrate the diminished killing of K562 cells by patients’ NK cells was due to a decrease in cytotoxic capacity rather than inefficient recognition or degranulation. This defect in cytotoxicity was reversible. Following 14 days in vitro culture with IL-2, NK cells from patient C08 killed K562 more efficiently than NK cells from HV11, their age matched counterpart (Figure 4G).

### T-cells are more differentiated in pediatric cancer patients

To determine if T-cells in pediatric cancer patients were altered in phenotype, manual gating of the concatenated fcs files from patients and controls was performed using CD45RA and CCR7 expression to divide CD4 and CD8 T-cells into naïve and memory subpopulations ^43^ (Figure 5A and 5C). Patients had lower frequencies of CCR7+ CD45RA+ naïve CD4 and CD8 T-cells with concomitant increases in CCR7+ CD45RA-central memory (CM), CCR7-CD45RA-effector memory (EM) and CCR7-CD45RA+ terminally differentiated effector memory (TEMRA) T-cells. Evaluating individual patients (Figure 5B and 5D) showed these changes in differentiation status were greatest for CD4+ T-cells; the frequency of T-cells across all four differentiation states were significantly different to those in age matched controls. Increased T-cell differentiation was not associated with prior cytomegalovirus infection (SI Fig 7).

**Figure 5.**
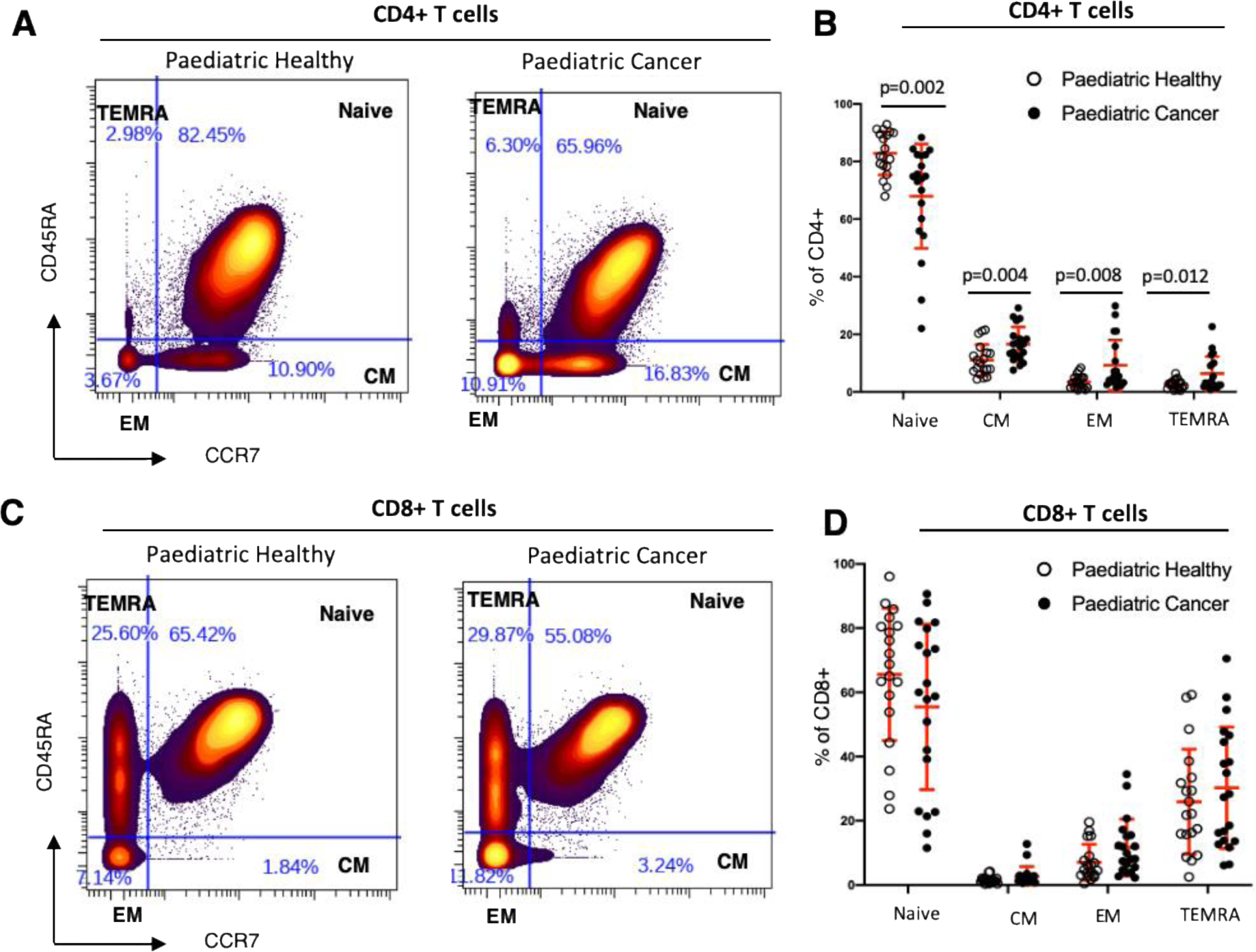
T cells are more differentiated in pediatric cancer patients. A) Results of manual gating CD4+ T-cells using fcs files concatenated from 19 healthy donors and 20 age matched controls. The percentage of each T-cell subset is shown as a percentage of total CD4 T cells. B) Result of gating shown in panel A for each individual. Unpaired t-test corrected for multiple comparisons with 5% FDR was used. The mean +/- 1 standard deviation is shown in red. C+D) The same analytical strategy for A and B, now investigating CD8+ T-cells.

To further evaluate differences in T-cell phenotype, a single t-SNE dimensionality reduction was performed on all CD3+ cells in the patient and control concatenated fcs files. There were marked differences between patients and controls as demonstrated by different spatial cell distributions in two-dimensional space (Figure 6A). To identify differences in the frequency of particular T-cell subsets, unsupervised FlowSOM clustering was used to assign T-cells into 40 meta-clusters (Figure 6B). Expression levels of key phenotypic markers on each meta-cluster are presented in Figure 6C as separate heatmaps for the 19 healthy control children (left panel) and 20 pediatric cancer patients (right panel) alongside fold difference of meta-cluster frequencies (middle panel). Visually comparing the two heatmaps, there was little difference in the marker expression for each meta-cluster apart from CXCR3, which was less intense across multiple meta-clusters in cancer patients. However, patients and controls had different frequencies of cells in many of the clusters. First, naïve CD4 and CD8 T-cell were present at higher frequency in healthy children. Second, EM and TEMRA CD4 and CD8 T-cell were present at higher frequency in patients. These changes confirm the earlier manual gated data (Figure 5).

**Figure 6.**
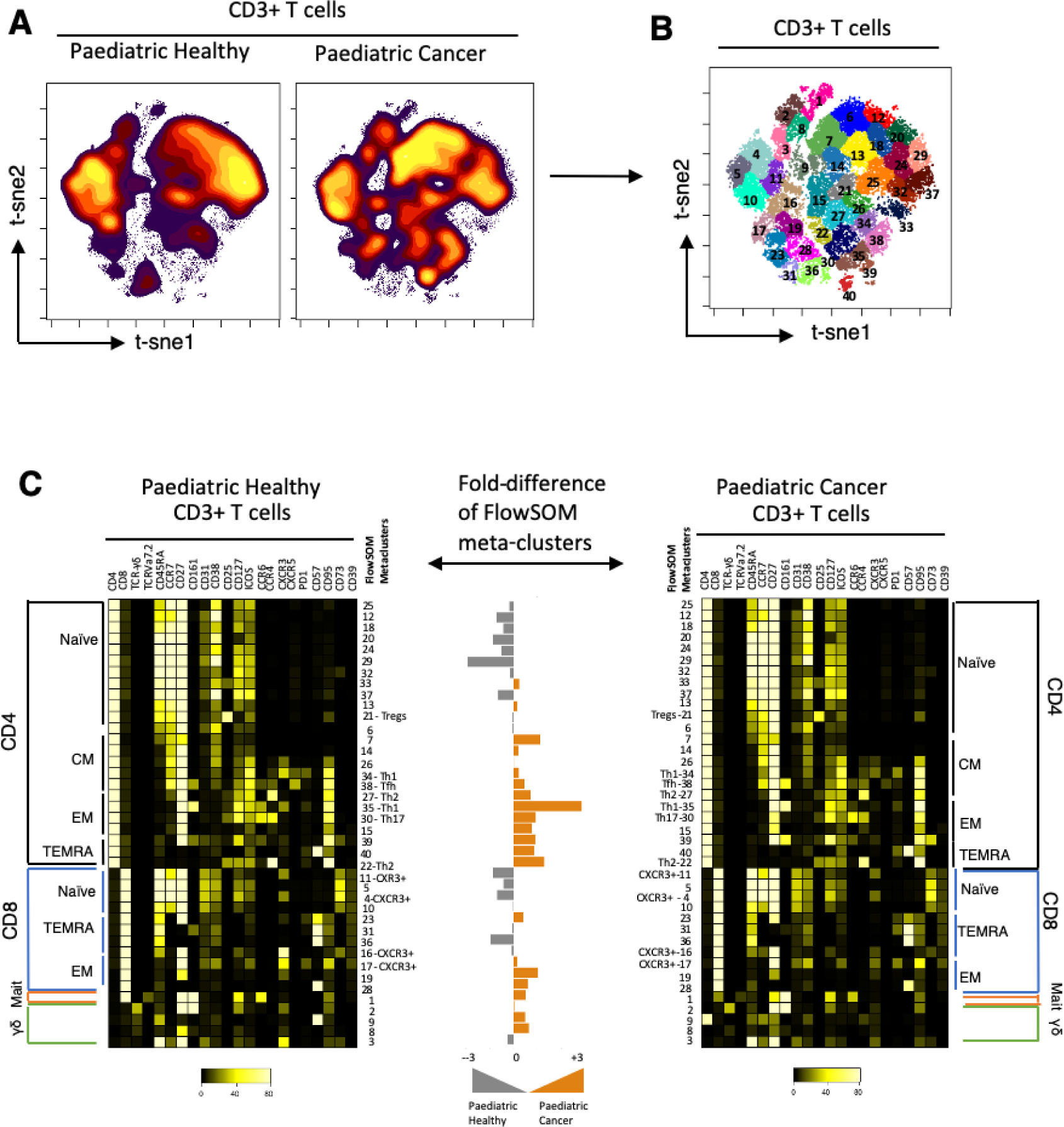
Unsupervised FlowSOM analysis of CD3+ T-cells reveals marked differences in the frequencies of T cell subsets in pediatric cancer patients relative to age matched controls. A) Results of tSNE dimensionality reduction performed on CD3+ cells from concatenated fcs files from patients and controls to assess differences on the spatial cell distributions in the two-dimensional space. B) Unsupervised clustering (FlowSOM) of T-cells was performed based on their spatial orientation on the tSNE axis, generating 40 meta-clusters. C) The expression of markers for each meta-cluster is shown on heatmaps generated for pediatric healthy donors (left panel) and pediatric cancer patients(right panel). The fold difference of the meta-clusters abundance between healthy donors and cancer patients is shown in the centre panel.

The unsupervised FlowSOM analysis provided additional insights. The largest difference in T-cell frequency between patients and controls was metacluster 35 (Th1 CD4+ T-cells based on CXCR3 expression) and metacluster 22 (Th2 CD4+ T-cells based on CCR4+ CCR6-). Frequencies of both were increased in patients. No difference in the abundance of metacluster 21 was observed (presumptive regulatory CD4+ T-cells, based on CD25^hi^CD127^lo^ expression). Manual gating of regulatory T-cells confirmed this result (SI Fig8). Regarding CD8+ T-cells, patients showed an increase in EM (meta-clusters 17, 19 and 28) and TEMRA (meta-cluster 23) cells. There was no difference in the frequency of CXCR3-positive CD8+ T-cells. Finally, patients had an increased frequency of gamma delta (γδ) T-cells (meta-clusters 8 and 9). Using a fluorescent cytometry panel (SI Table 5), we found this increase was solely due to effector Vδ1 γδ T-cell cells (p=0.0126; SI Fig 9) a subset with adaptive immunobiological properties ^44^. The innate-like Vδ2 γδ T cells were of similar frequency in patients and controls although there was a clear difference in phenotype, with levels of perforin lower in patients’ cells (p=0.012, SI Fig 9).

### Pediatric cancer associated immune perturbations vary by age

Finally, because the immune system develops throughout childhood we investigated if the immune changes observed in cancer patients were influenced by age. Applying robust linear regression modelling to the mass cytometry data showed there was little effect of age on the total frequency of CD4 and CD8 T-cells and monocytes (Figure 7A). Although the frequency of T-cells did not vary with age there was, however, a striking age-related effect on the differentiation status of CD4 and CD8 T-cells in cancer patients (Figure 7B). With increasing age at diagnosis the proportion of naïve T-cells decreased and the proportion of effector memory and TEMRA T-cells increased to levels seen in healthy adults (shown as box and whisker plots, n=19 adults, median age=54 range=31-79 years).

**Figure 7.**
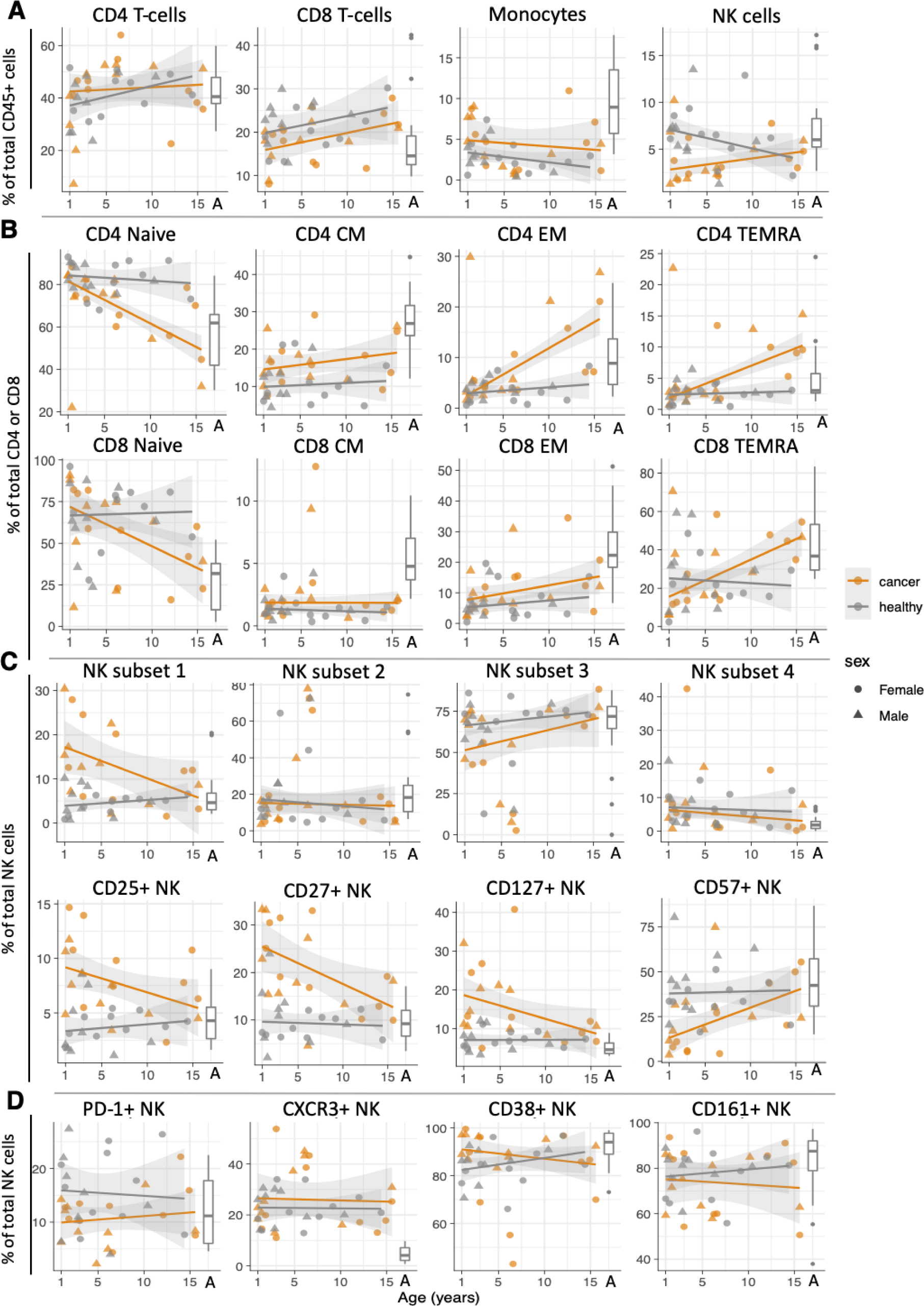
Influence of age on immune cell frequency and phenotype. The frequencies of each cell population as a percentage of A) CD45+ cells, B)CD4 (upper row) or CD8 (lower row) T cells or C) total NK cells. Orange symbols show cell frequencies for cancer patients, grey symbols show cell frequencies for healthy donors. The regression line of a robust linear model with 95% confidence intervals applied to each cell subset in patients or healthy donors is also shown. The box and whiskers plots shows the frequency of same cell subsets in healthy adults (n=19, age range= 31-75 years, median age=54 years)

In contrast to other immune cells, the frequency of total NK cells was age dependent, being lower in younger, but not older, children with cancer (Figure 7A, right panel). We also identified a striking effect of age on NK differentiation (Figure 7C). Using the CD16 and CD56 schema of NK cell development we demonstrated that the proportion of immature CD56^bright^ CD16^-^ subset 1 NK cells was increased in young (<8 years old), but not older, cancer patients. In addition, other markers of NK immaturity, including CD25, CD27 and CD127 were also increased in young patients. In accordance with these results, the proportion of NK cells positive for CD57, which marks differentiated NK cells capable of cytotoxicity, was low in younger children with cancer and increased with age. This age-related effect did not apply to other NK phenotypic markers such as CXCR3, CD38, CD161 and PD1, though we noted the latter was always expressed on a smaller proportion of cancer patient’s NK cells (figure 7D).

Collectively, our results show that young children with cancer have a decreased frequency of peripheral NK cells and that these NK cells exhibit increased immaturity. These perturbations were not seen in older children with cancer who instead had increased frequencies of effector and TEMRA CD4 and CD8 T-cells. These age-related changes were observed only in cancer patients and not in healthy children.

## Discussion

Pediatric cancers occur between the ages of 1 and 16 years, a time when the immune system is undergoing major developmental changes as it progresses from a neonatal to an adult state^1^. Surprisingly little is known about the immune system of normal children in this age range. Cancer is known to alter systemic immunity and whilst this has been extensively explored in association with adult cancers there is very little knowledge of the immune status of pediatric patients. By performing a systematic single cell high dimensional analysis, we have generated baseline data on normal cell-mediated immunity in children and have identified notable differences in systemic immunity in pediatric cancer patients. In the younger (< 8 years) patients the frequency, phenotype and function of NK cells was reduced, whereas in the older patients (> 8 years) the T cell differentiation profile was altered.

Most pediatric cancers arise in young children 1-5 years old ^1^. NK cells are likely to be critical anti-tumor effectors in this age range because: i) the T-cell system is immature and Th2 skewed ^6^ and ii) the embryonal tumors that are more common in this age range lack HLA-I expression ^15–18^ but can still be killed by NK cells ^15,19^.These observations raise an interesting paradox: how can nascent NK-sensitive embryonal tumors develop, proliferate and persist in children? Our data shows NK function in the periphery of young children with cancer is compromised at multiple levels. We observed that the frequency of NK cells in the periphery is decreased. This could be due to migration of NK cells from the periphery into tumors, however, arguments against this are: i) there is no difference in the proportion of NK cells in the periphery expressing the CXCR3 receptor essential for tumor entry and ii) pediatric tumors generally contain few immune cells ^45^. An alternative explanation, given that the lifespan of NK cells is only two weeks, is that NK cell homeostasis is disturbed. This hypothesis is supported by our observation that the phenotype of the low frequency NK cells present in pediatric patients is substantially altered, as demonstrated by the altered proportions of immature and mature NK cells.

In adult cancer patients, tumor cells have been shown to secrete NK ligands that bind to NK cell receptors abrogating their function. We explored whether this mechanism might also occur in pediatric patients, focusing on sULBP2 and sMICA based on a report that plasma levels of both were increased in young neuroblastoma patients ^46^. We detected both ligands in our patients’ plasma but concentrations were lower than those in healthy age-matched children. Others have reported soluble NK ligands can be readily detected in healthy children’s plasma ^47^. We note that the neuroblastoma study ^46^ did not report the age of their healthy donor comparator group; we therefore suggest an age mismatch between the patient and control groups in their study may account for our different results. If soluble NK ligands were indeed inhibiting NK function in pediatric patients then we would have expected to have seen lower degranulation and cytokine production by NK cells from patients compared to their age-matched healthy counterparts. This was not the case, arguing against an inhibitory role for soluble NK ligands in pediatric cancer.

Lysis of K562 cells by NK cells from four of the five patients tested was substantially reduced. Although NK cells from patients expressed higher levels of the inhibitory receptor NKG2A, this cannot explain their decreased function because i) K562 cells lack the HLA-E molecule that inhibits NK function via NKG2A ^48^ and ii) as noted above there was no decrease in cytokine production and degranulation by these patients’ NK cells. We also found levels of the activatory receptor NKG2D, through which K562 is targeted by NK cells ^49^, were equivalent. Therefore, the decrease in NK cytotoxicity observed for patients is likely due to the lower levels of perforin and granzyme-B in their NK cells and the decreased proportion of mature NK cells, particularly CD57+ cells that are the strongest mediators of cytotoxicity. We cannot rule out additional contributions by alterations in other activatory and inhibitory NK receptors which were not measured in our study; a detailed characterization of NK cells is therefore a priority for future work. The observation that the one patient with normal NK cytotoxicity had nehproblastomatosis, a pre-malignant disease, rather than cancer is intriguing. With the caveat that it is only one patient, it suggests compromised NK cytotoxicity may not occur until later in the disease process.

For children over eight years of age in our patient cohort we did not detect any changes in NK frequency or phenotype compared to their age matched controls. Instead, we saw substantial changes in T-cell differentiation for both the CD4+ and CD8+ subsets, with the proportion of naive cells decreased and concomitant increases in the proportion of effector memory and TEMRA T-cells. The magnitude of these changes in differentiation status, relative to age matched healthy children, were striking with frequencies of TEMRA in older children resembling those in healthy middle-aged adults. Although we did not measure the functional capacity of TEMRA cells in cancer patients, they were largely negative for CD57 (Si Fig 8), a marker of terminally differentiated T-cells that have short telomeres, high apoptosis sensitivity and replicative senescence ^50^. These expanded memory and TEMRA T-cells are therefore likely to be functional. This raises the question whether the expanded memory T-cells could be specific for tumor antigens. Increases in highly differentiated TEMRA cells occurs in several pathologies where they could be antigen driven. These include transplantation, where TEMRA cells are clonally expanded and increased frequencies are associated with tissue rejection ^51^; lupus and long-term infection with cytomegalovirus which results in large virus specific TEMRA cell populations in the elderly ^52^. It is notable that in adult cancer patients, increased TEMRA are associated with better prognosis following anti-PD1 immunotherapy ^53^. In the context of our study, where patients were receiving conventional cytotoxic therapy and not immunotherapy, this potential association could not be explored. However, it should be considered for on-going immune-oncology trials in children with cancer. Given that T-cell differentiation is simple to measure using fluorescent flow cytometry, this question could also be tackled in children receiving current standard of care treatments by implementing cytometry into routine care. Emerging reports show checkpoint inhibitors elicit poor responses in pediatric patients ^4^. This may be due to the low mutational burden creating fewer neoantigens ^14^. Our results provide an additional, but not necessarily mutually exclusive, explanation why this may be the case. Our results also provide preliminary insights into a rational basis for improving cancer immunotherapy in children. First, our data show that unlike adult cancer patients ^54^, children with cancer do not have increased regulatory T-cells (SI Fig 9). Therefore, targeting these cells in children may not augment anti-tumor immunity. Second, assuming at least a proportion of the increased effector T-cells we observed in older children are tumor specific, identifying mechanisms to increase their access into tumors and blocking relevant inhibitory pathways may be beneficial. Third, in young children, particularly those with embryonal tumors, restoring NK cell function should be a priority. We demonstrate reversal of NK dysfunction though co-culture with interleukin-2 but other NK-modulating therapeutic approaches are already being explored in adult patients. These include infusion of interleukin-15, a cytokine pivotal for NK development in vivo, the use of an IL-15 agonist antibody or an anti-NKG2A blocking antibody ^55,56^. Patients with high risk neuroblastoma currently receive a monoclonal antibody specific for the tumor protein GD2 which is thought to act through NK cell mediated antibody dependent cellular cytotoxicity ^57^. Although anti-GD2 has been combined with NK infusion in small trials, these were not designed to measure benefit from NK addition^58^. Our data showing decreased NK frequency, maturity and cytotoxicity in young patients, in whom neuroblastoma is prevalent, suggest NK cell augmentation should be considered as a priority for use alongside anti-GD2.

Our patient cohort included a range of different solid tumour types and within this limited sample set our observations did not appear to be disease specific. Further exploration of a larger cohort of cancer patients to examine potential disease-specific changes in systemic immunity, in addition to the age specific changes we identified, is warranted. In summary, high dimensional single cell analysis of pediatric cancer patients revealed important changes in NK and T-cell subsets influenced by age. Our results provide a rational basis for selecting effective immunotherapies for pediatric cancer patients and strategies for future trials.

## Material and Methods

### Study design and patient recruitment

All samples were obtained from children at Birmingham Women’s and Children’s Hospital as part of an ethically approved study (TrICICL) in accordance with the declaration of Helsinski and approved by South of Birmingham Research Ethics Committee (IRAS: 233593). Written informed consent was obtained from all participants or their legal guardians. Samples were collected and stored at room temperature and processed within 24 hours.

### Blood Samples and PBMC Isolation

Peripheral blood samples from pediatric cancer and healthy patients were collected in EDTA tubes and peripheral blood mononuclear cells (PBMCs) were isolated from blood samples using SepMate tubes (Stemcell technology) as per the manufacturer’s protocol. PBMCs were cryopreserved in medium containing DMSO^59^ before transfer to liquid nitrogen for long-term storage.

### Mass Cytometry (CyTOF) panel

Two panels of antibodies were designed and used for deep immune phenotypic analysis (SI Tables 1 and 3). Antibodies were purchased pre-conjugated from Fluidigm or unconjugated from Biolegend and conjugated in house using the Maxpar antibody labelling kit from Fluidigm following manufacturer’s protocol.

### Mass Cytometry (CyTOF) staining and analysis

Details of the staining protocol is provided in supplementary methods. Bead normalised data were analysed in Cytobank. Samples were debarcoded, down sampled to 4751 cells per sample and all fcs files from patients or healthy donors concatenated for further analysis. Gating strategies are provided in supplementary Figure 10. Dimensionality reduction and clustering were performed in Cytobank using the ViSNE implementation of tSNE and FlowSOM respectively. UMAP was performed using the UWOT package and robust linear modelling performed using the MASS package in R version 3.5.3. Marker enrichment modelling v3.0 was performed in R using code downloaded from GitHub. Additional statistical tests, multiple t-test (correcting for multiple comparisons using the Benjamini, Krieger and Yekuteli two stage step up method with 5% FDR) or unpaired Mann-Whitney test, were performed in GraphPad prism 8.

### NK functional assays and phenotype

NK cell cytotoxicity, degranulation and cytokine production, were evaluated by co-culturing with K562 Cells (Erythroleukemia cell line) in vitro (see supplementary methods). An intracellular flow cytometry panel was used to measure perforin and granzyme-B levels (SI Table 4). Data were acquired on an LSR-II cytometer and analyzed in Cytobank.

## Acknowledgements

We thank the research nurses at Birmingham Women and Children’s Hospital Sara-Jane Stanley, Jane Cooper and Cay Shakespeare for their help and support recruiting to the TRICICL study. We thank the patients, their families and all healthy volunteers for their participation in the study.

## Authors contributions

ES, PK and GT conceived and designed the study. Samples were collected and processed by ES, TH, NM and GT. Data were generated by ES, NK, CW and GT. Data analysis and interpretation were performed by ES, JI, SB, CW, BW, JZ, PM, PK and GT. Manuscript writing was performed by ES, JI, PK and GT. The final version of the manuscript was approved by all authors who are all accountable for the work.

## Data and materials availability

Mass cytometry data are available for download from flow repository.

## Supplementary methods

### Mass Cytometry staining protocol

Cryopreserved PBMC were recovered into cell culture media (RPMI, 10% Foetal Bovine Serum, penicillin 50U/ml and streptomycin 50U/ml) and washed once. Equal numbers of viable cells were transferred into FACS tubes for barcoding with CD45 specific antibodies conjugated with one of four different metals. PBMCs from cancer patients, pediatric healthy donors and adult volunteers were barcoded using a batch randomisation scheme to avoid bias. Also included in the experiments were buffy coat PBMCs, separately barcoded with each of the four different metals and stained in parallel to assess technical variability. CD45-specific antibodies were labelled with: 89Y (Yttrium), 114 Cadmium (Qdot605), 115 Indium or Platinum. After 20 minutes incubation, cells were washed and combined together for further staining.

A master-mix of all 36 phenotyping antibodies was prepared by adding the appropriate pre-tested dilutions into filtered cell stain media (CSM - phosphate buffered saline + 0.5% Foetal Bovine Serum + 0.02% sodium azide). After 30 minutes incubation, live dead rhodium stain (Fluidigm) was added. The samples were washed twice then fixed overnight with freshly prepared 1.6% formaldehyde. The following day, cells were incubated with iridium intercalator (Fluidigm) solution for one hour and analysed on a Fluidigm Helios mass cytometer using an acquisition rate less than 500 events per second. Immediately prior to acquisition cells were washed once in CSM and twice in deionised water then filtered through a 70μm cell strainer. Four element calibration beads (Fluidigm) were used for data normalisation using the Helios data acquisition software. Normalised data were uploaded to Cytobank for further analysis including de-barcoding, manual and automated analysis.

### NK cytotoxicity assay

Cytotoxicity of NK cells was determined by measuring their ability to kill the erythroleukemia cell line K562. PBMCs were recovered and incubated overnight in culture media (RPMI, 10% Foetal Bovine Serum, Penicillin 50U/ ml and streptomycin 50U/ml) plus 200IU/ml IL-2 (Peprotech). Cells from each donor were co-cultured with CFSE labelled K562 (CFSE tracking kit, Biolegend) correcting for differences in NK frequency in PBMCs so the effector to target ratios were defined. Cells were cultured at 37°C in cell culture media (RPMI, 10% foetal bovine serum, penicillin 50U/ ml and streptomycin 50U/ml). The following day, cells were recovered, washed once with PBS and incubated for 10 minutes with efluor 780 live dead stain (ebioscience). Counting beads (ebioscience) were added to the sample and data acquired using an LSR-II flow cytometer. Cytotoxicity was calculated using the formula cytotoxicity= [(expected live target cells - live target cells)/ expected live target cells] x100.

### NK cell degranulation assay

NK cell degranulation was measured similarly to NK cytotoxicity. PBMCs, recovered and incubated overnight in IL-2 containing media as described above, were co-cultured with K562 cells at defined NK:K562 cell ratios. Anti-CD107a-FITC antibody (Biolegend) was added at the beginning of the assay at 1ul/well. After one hour, monensin and brefeldin A (Biolegend) were added to the assay as per manufacturer’s protocol. At the end of the incubation cells were stained with fluorophore-conjugated antibodies specific for CD3, CD16, CD56 and CD57. Cells were then fixed and permeabilized using the eBioscience fixation and permeabilization kit and intracellular staining for INF-g and TNF-a performed. Samples were then acquired on an LSR-II flow cytometer and data analysed in Cytobank.

**SI Table 1.**
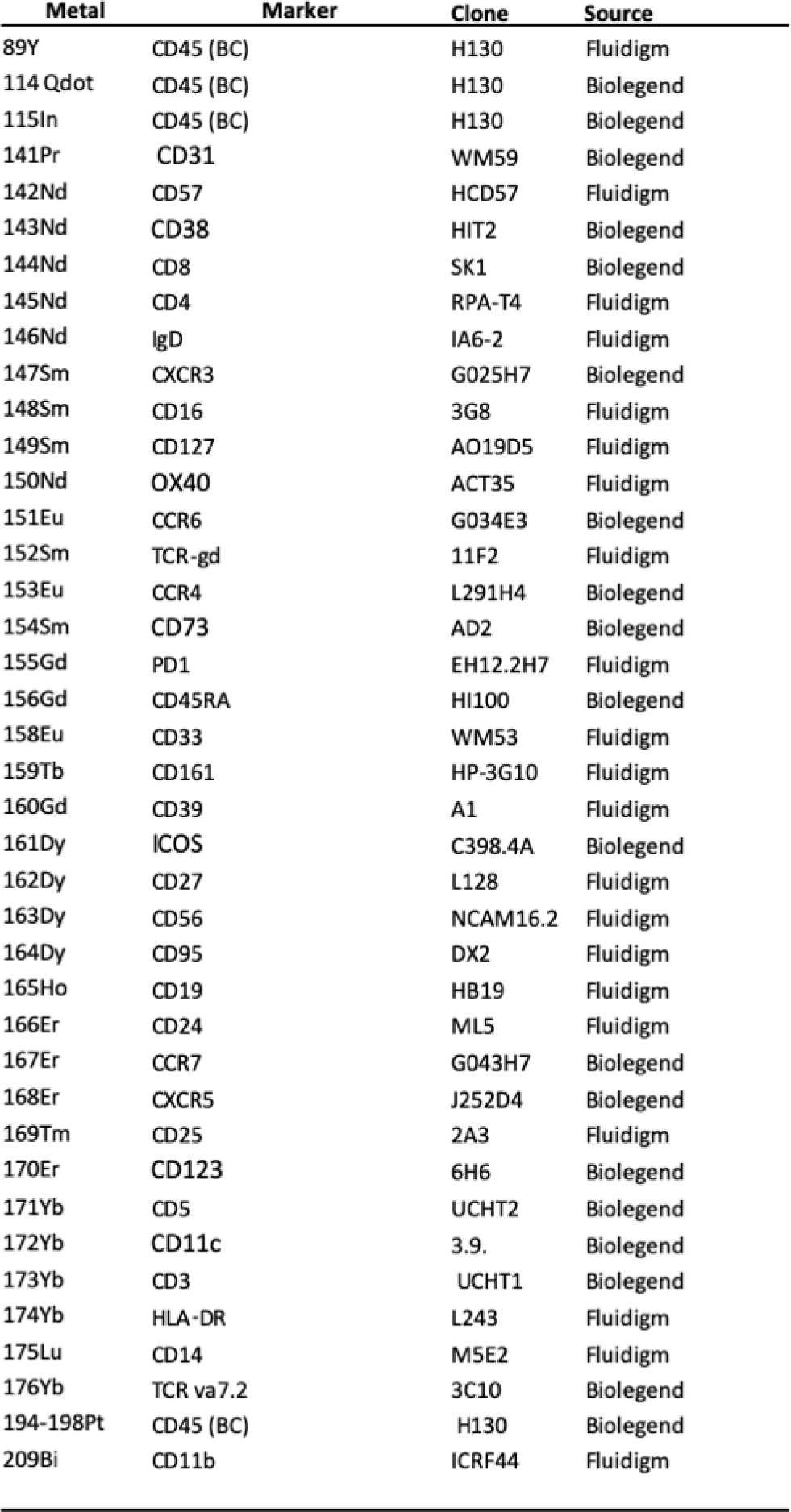
Mass cytometry antibody panel used to characterise all donors recruited to the study.

**SI Table 2.**
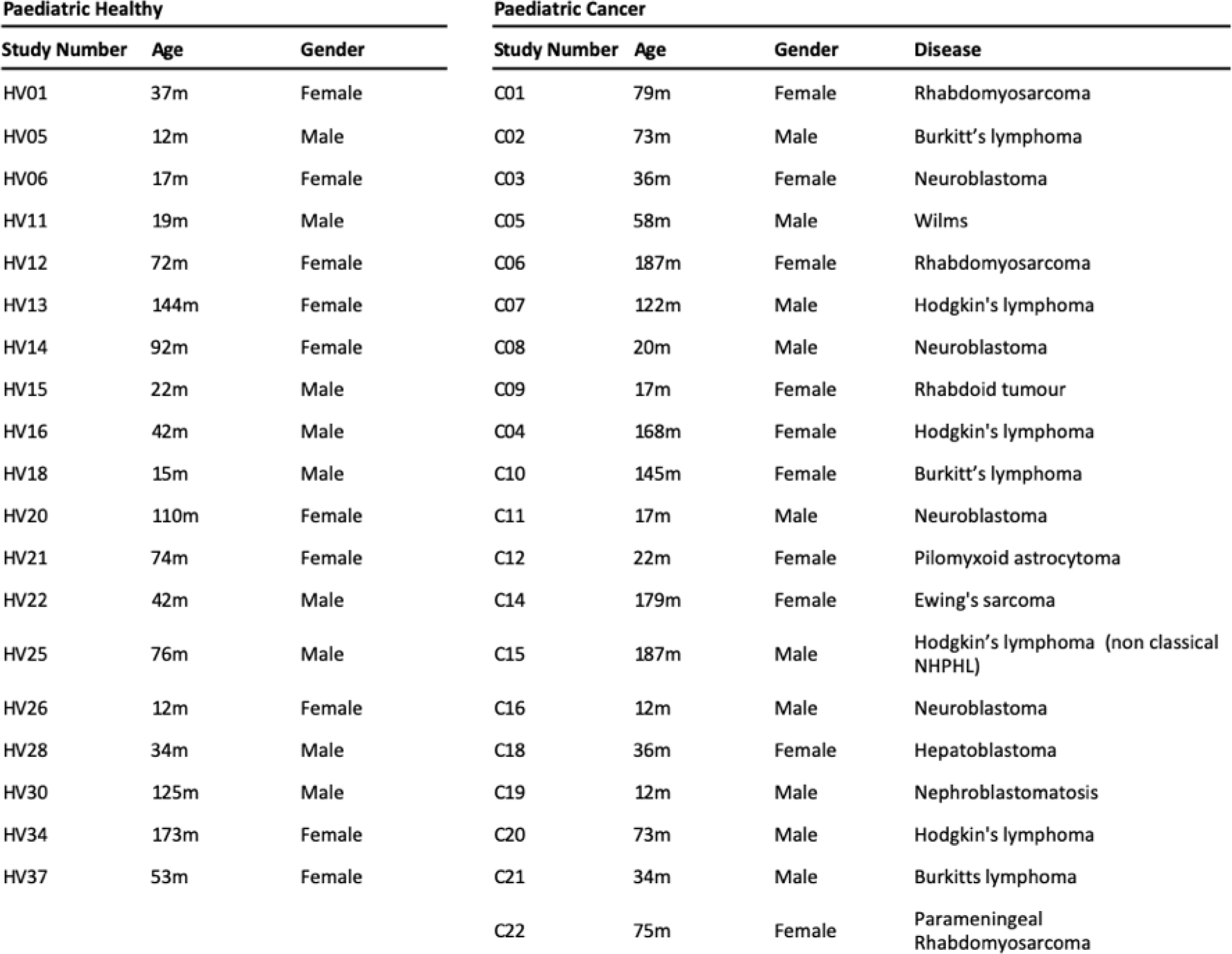
Demographic data for the paediatric healthy donors and paediatric cancer patients. The diagnosis for each patient is also shown. Age is provided in months.

**SI Table 3.** Mass cytometry antibody panel used to characterise NK cell phenotype

| Metal | Marker | Clone | Source |
| --- | --- | --- | --- |
| 89Y |  |  |  |
| 114 Qdot |  |  |  |
| 115In | CD45 (BC) | H130 | Biolegend |
| 141Pr | gamma-delta 2 | 123R3 | Milteny |
| 142Nd | CD57 | HCD57 | Fluidigm |
| 143Nd |  |  |  |
| 144Nd | CD8 | SK1 | Biolegend |
| 145Nd | CD4 | RPA-T4 | Fluidigm |
| 146Nd |  |  |  |
| 147Sm | CXCR3 | G025H7 | Biolegend |
| 148Sm | CD16 | 3G8 | Fluidigm |
| 149Sm | CD127 | AO19D5 | Fluidigm |
| 150Nd |  |  |  |
| 151Eu | CCR6 | G034E3 | Biolegend |
| 152Sm | CD27 | M-T271 |  |
| 153Eu | CX3CR1 | 2A91 | Biolegend |
| 154Sm | TIGIT | A15153G | Biolegend |
| 155Gd | PD1 | EH12.2H7 | Fluidigm |
| 156Gd | CD45RA | HI100 | Biolegend |
| 158Eu |  |  |  |
| 159Tb | Nkp30 | Z25 | Fluidigm |
| 160Gd | gamma-delta 1 | REA173 | Milteny |
| 161Dy |  |  |  |
| 162Dy | Nkp46 | BAB281 | Fluidigm |
| 163Dy | CD56 | NCAM16.2 |  |
| 164Dy | CD95 | DX2 |  |
| 165Ho | 1° NKG2C-PE<br>2° Anti-PE | 134591,<br>PE2 | R&D Biosystems,<br>Fluidigm |
| 166Er | NKG2D | ON72 | Fluidigm |
| 167Er | CCR7 | G043H7 | Biolegend |
| 168Er | CXCR5 | J252D4 | Biolegend |
| 169Tm | NKG2A | Z199 | Fluidigm |
| 170Er | CD161 | HP3610 | Biolegend |
| 171Yb | DNAM | DX11 | Fluidigm |
| 172Yb |  |  |  |
| 173Yb | CD3 | UCHT1 | Biolegend |
| 174Yb |  |  |  |
| 175Lu | CD14 | M5E2 | Fluidigm |
| 176Yb | TCR-va 7.2 | 3C10 | Biolegend |
| 194Pt | CD45 (BC) | H130 | Biolegend |
| 198Pt | CD45 (BC) | H130 | Biolegend |
| 209Bi | CD11b | ICRF44 | Fluidigm |

**SI Table 4.** *e-Fluor 780 live dead stain detected on APC-CY7 detector. Fluorescent flow cytometry antibody panel used for further NK cell characterisation

| Fluorochrome | Antibody | Clone | Source |
| --- | --- | --- | --- |
| e450 | CD3 | SK7 | eBioscience |
| APC | CD56 | CMSSB | eBioscience |
| PeCyC7 | CD16 | eBioCB16 | eBioscience |
| Percpcy5.5 | CD57 | QA17A04 | Biolegend |
| FITC | CD107a | H4A3 | Biolegend |
| AF700 | TNF-a | Mab11 | Biolegend |
| PE | INF-g | 4S.B3 | Biolegend |
| APC-Cy7 | CD19 | HIB19 | eBioscience |
| APC-Cy7 | CD14 | 6ID3 | eBioscience |
| APC-Cy7* | Live-Dead |  | Biolegend |
| PeCy7 | perforin | dG9 | eBioscience |
| PE-texas red | granzyme b | GB11 | Invitrogen |

**SI Table 5.** Fluorescent flow cytometry antibody panel used for γδ T-cell characterisation

**SI Table5**
| Fluorochrome | Antibody | Clone | Source |
| --- | --- | --- | --- |
| BV510 | live/dead | Zombie Aqua | BioLegend |
| BV421 | CD3 | UCHT1 | BioLegend |
| BV650 | CD8 | SK1 | BD Biosciences |
| FITC | Vd1 | REA173 | Miltenyi |
| PE | Vd2 | 123R3 | Miltenyi |
| PE-Cy5 | Vg9 | IMMU 360 | Beckman Coulter |
| PE Dazzle 594 | CD27 | M-T271 | BioLegend |
| BV711 | CD45RA | HI100 | BD Biosciences |
| BV605 | CD45RO | UCHL1 | BioLegend |
| APC Fire 750 | CD161 | HP-3G10 | BioLegend |
| AF647 | Hobit | Sanquin-Hobit/1 | BD Biosciences |
| PE-Cy7 | Perforin | B-D48 | BioLegend |
| BV786 | Tbet | O4-46 | BD Biosciences |

**SI Fig 1.**
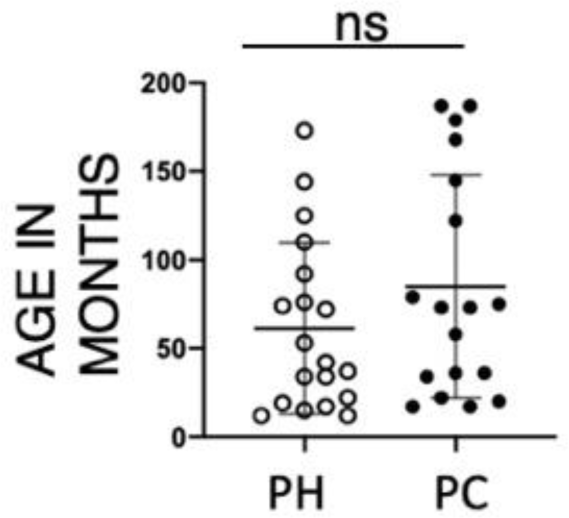
Age distribution of paediatric healthy (PH) or paediatric cancer (PC) donors. Error bars show mean +/-1 standard deviation. There was no significant difference in age between the two groups (unpaired t-test).

**SI Fig 2.**
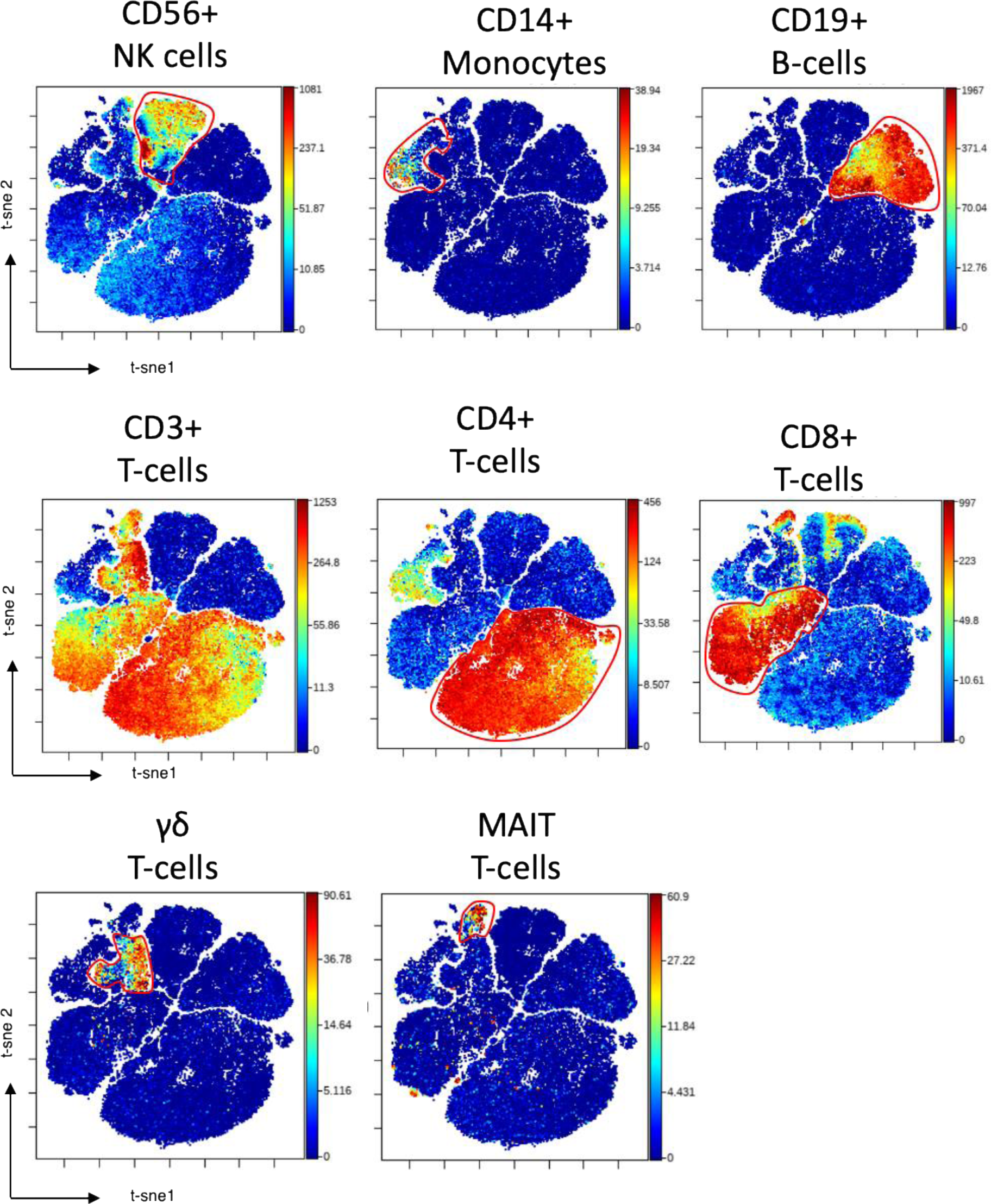
Gating strategy used to separate total immune cells into subsets for analysis (Fig 1-4 and 6-8). Each subset is determined by the indicated marker. Intensity of staining is represented by colour, with levels shown on the right hand side of each plot.

**SI Fig 3.**
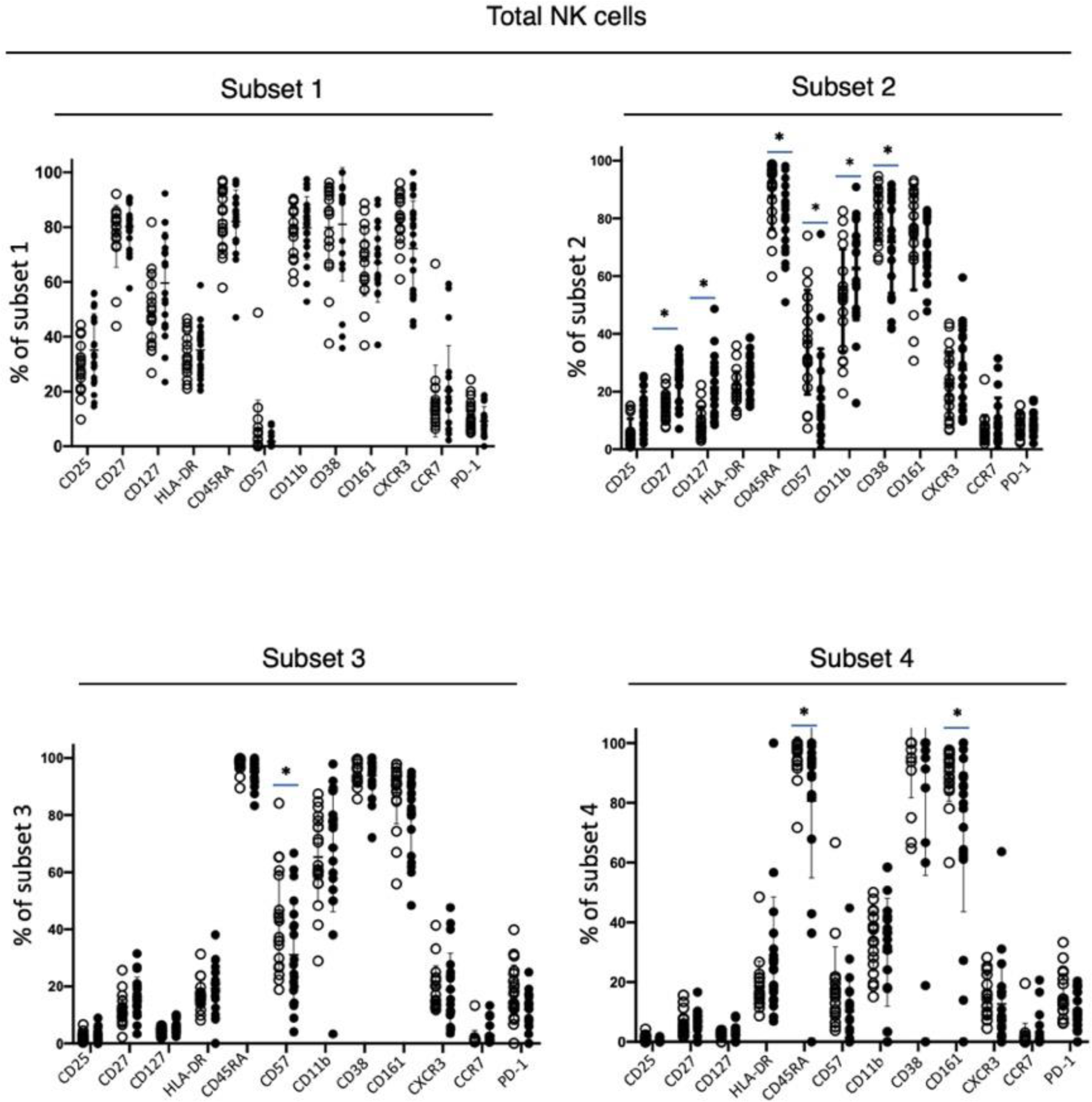
Frequency of NK cells within each of the four NK subsets (defined using CD16 and CD56) showing the percentage of cells positive for each of the indicated markers. The error bars show the mean +/- one standard deviation. Statistical significance testing was performed by unpaired t-test correcting for multiple comparison with 5% FDR. Statistical significant results are indicated by an asterisk.

**SI Fig 4.**
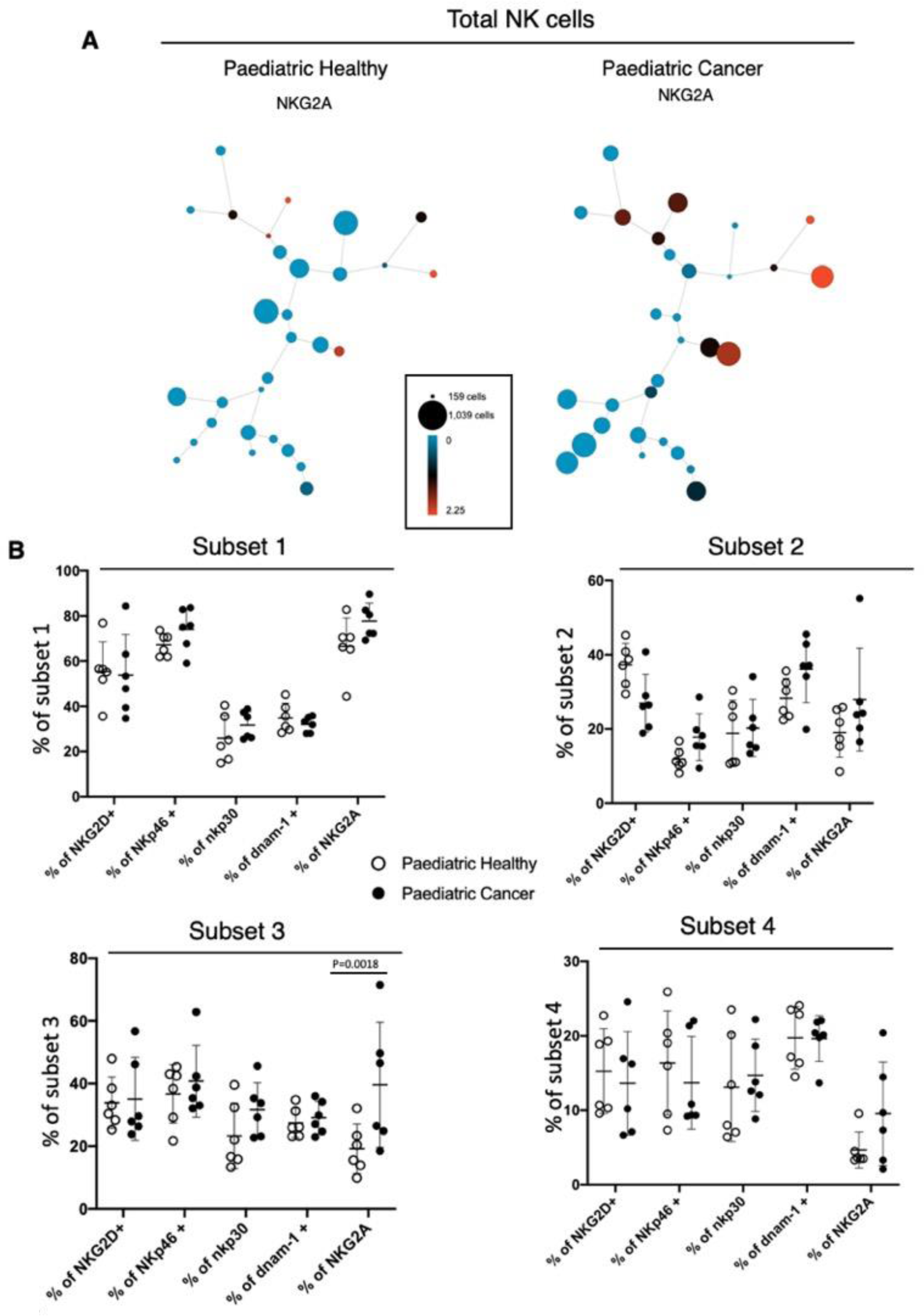
A) Total NK cells were clustered using the SPADE algorithm in Cytobank. Spade nodes are coloured by the level of NKG2A expression. Size of each node reflects the numbers of cells. B) Frequency of NK cells within each of four NK cell subsets (defined using CD16 and CD56) showing the percentage of cells positive for each NK receptor. The error bars show the mean +/- one SD. Unpaired t-tests, correcting for multiple comparisons with 5% FDR, were performed with significant results indicated by an asterisk.

**SI Fig 5.**
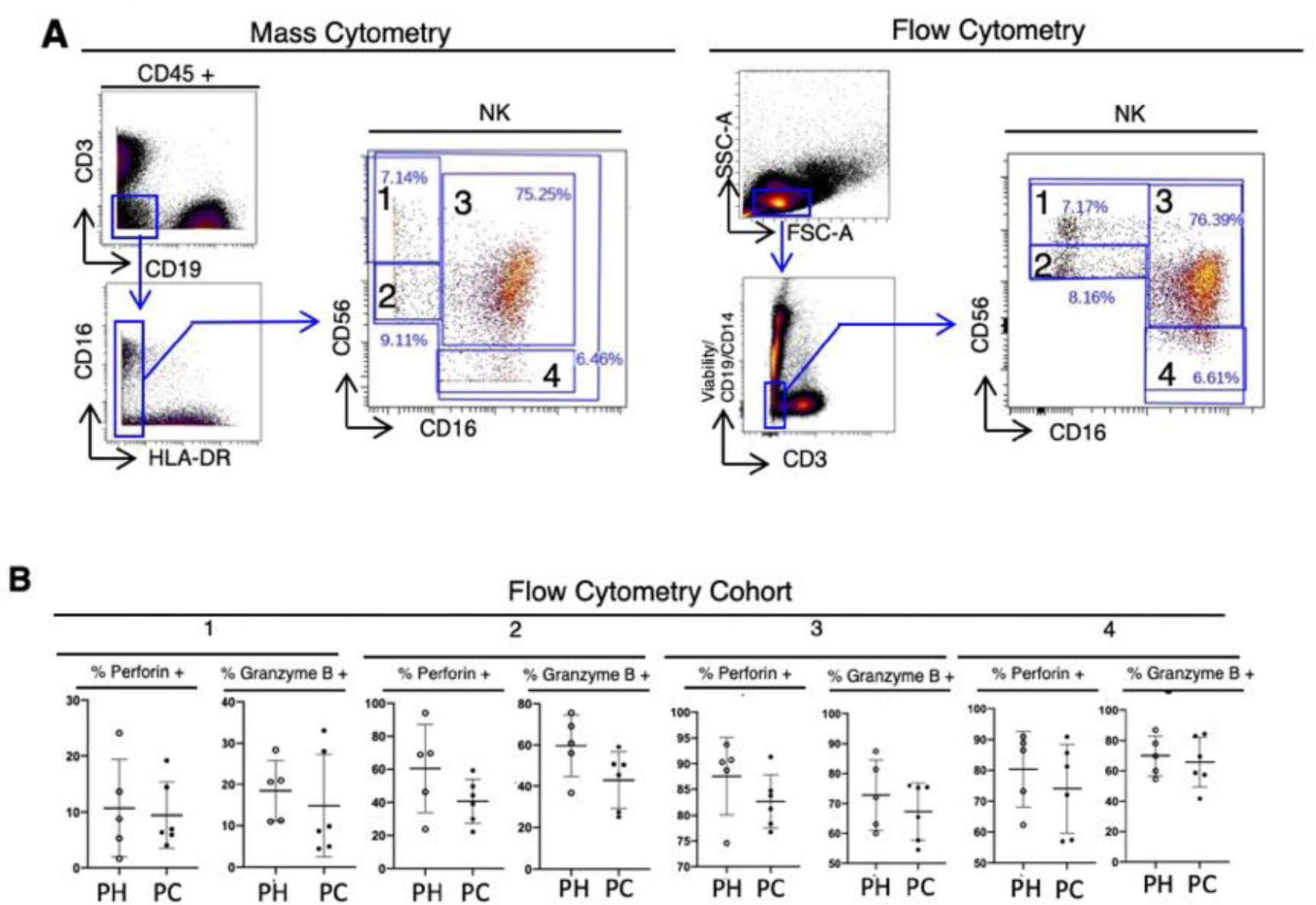
A) Gating strategy used to investigate the four NK cell subsets (defined using CD16 and CD56) from mass and flow cytometry data. B) Frequency of NK cells within each NK subset showing the percentage of cells positive for perforin or Granzyme B. The error bars show the mean +/- one SD. Unpaired t-test, correcting for multiple comparisons with 5% FDR, were performed with no significant results identified.

**SI Fig 6.**
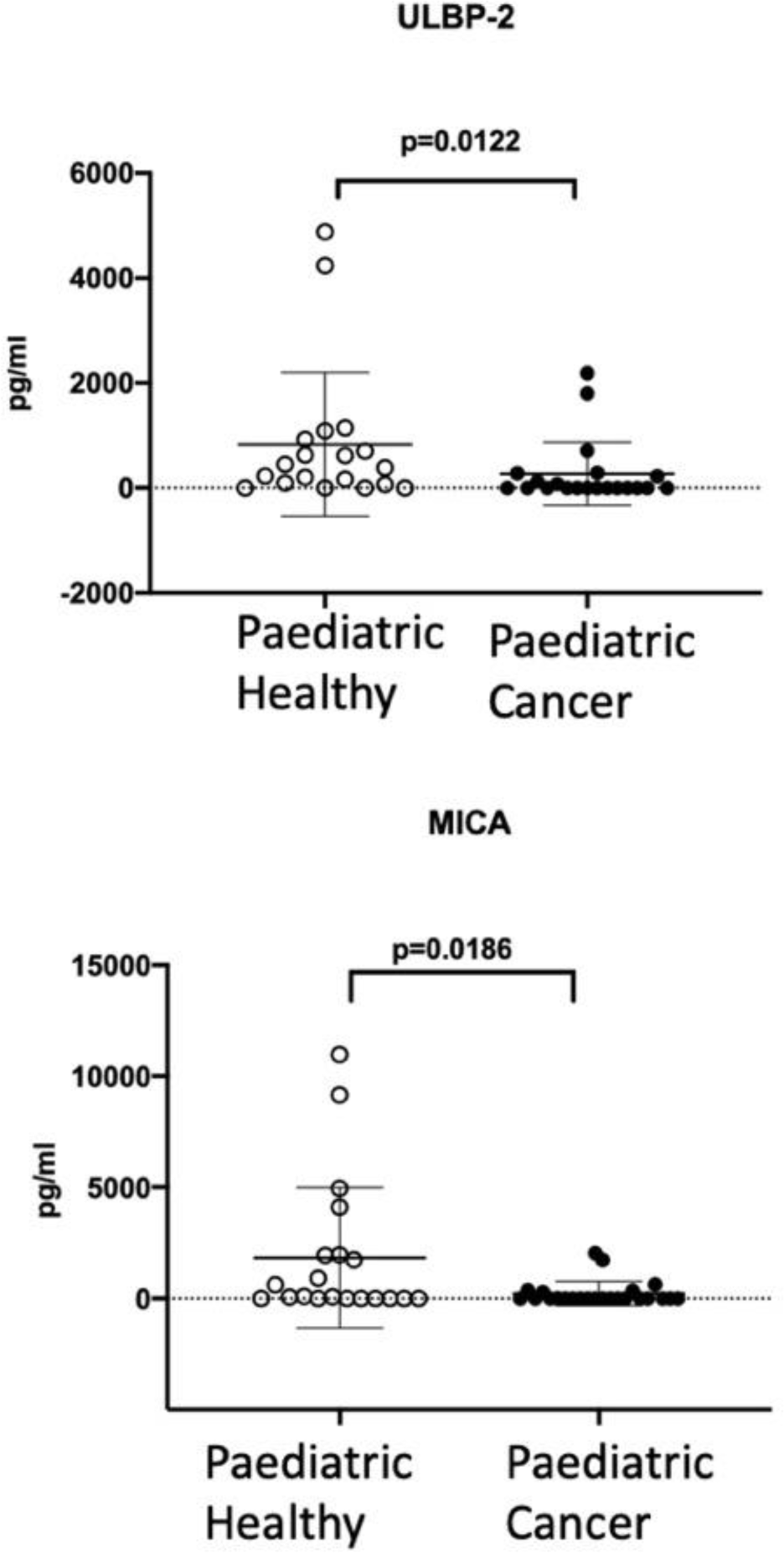
Results of ELISAs measuring NKG2D ligands ULBP-2 or MICA. The error bars show the mean +/- one SD. Mann-Whitney tests were used to a compare paediatric healthy and paediatric cancer donors (ULBP-2, p=0.0122; MICA, p=0.0186).

**SI Fig 7.**
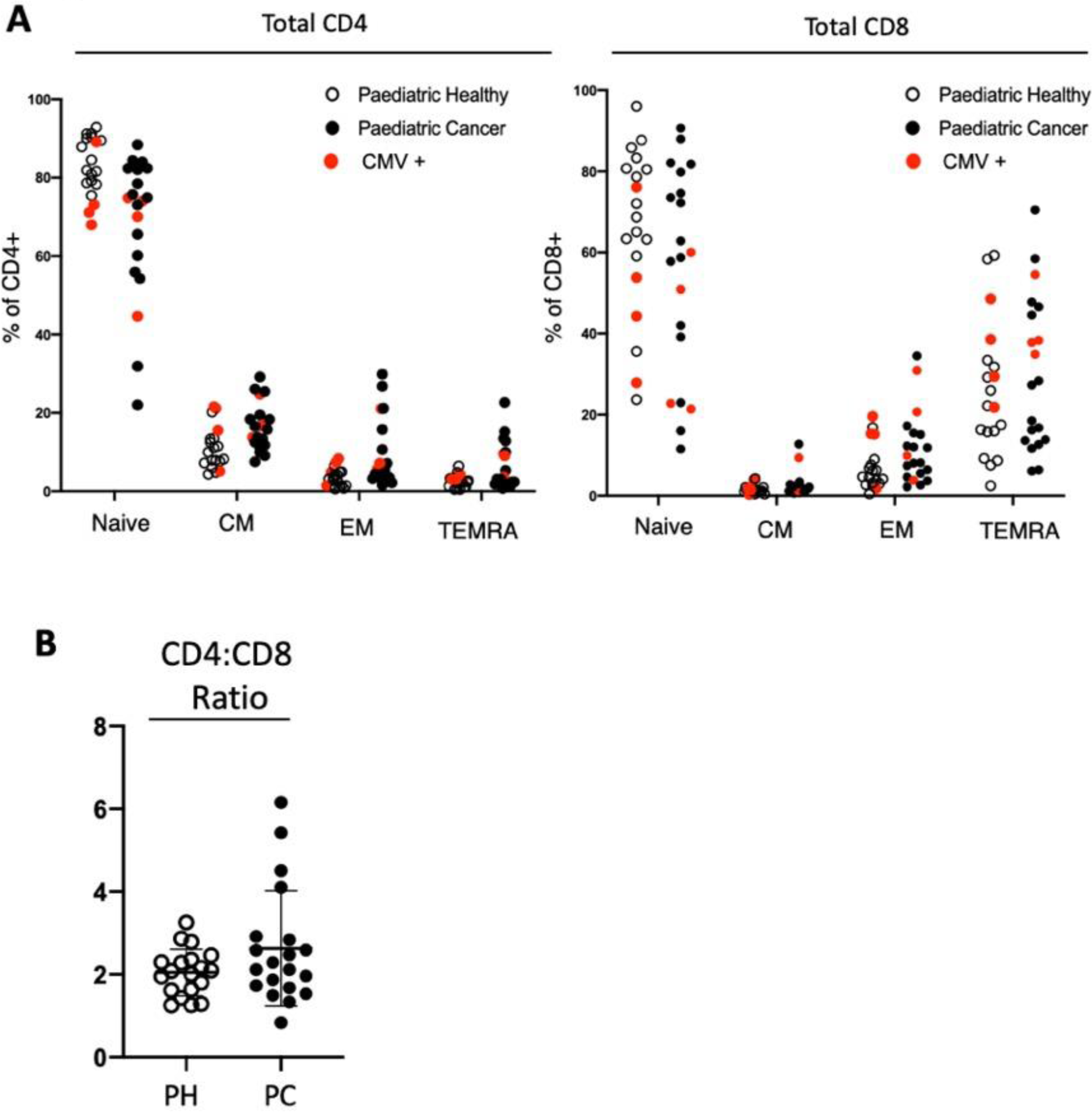
A) The frequency of CD4+ and CD8+ T cells subsets (Naïve; CM, Central Memory; EM, Effector Memory and TEMRA, T effector memory CD45RA revertant) is shown for individuals as a percentage of total CD4+ or CD8+ T cells. Donors that tested positive for cytomegalovirus IgM or IgG antibodies are coloured red. B) CD4:CD8 T-cell ratio for healthy donors and cancer patients are shown for individuals. The error bars show the mean +/- one SD. Mann-Whitney test was performed with no significant difference detected.

**SI Fig 8.**
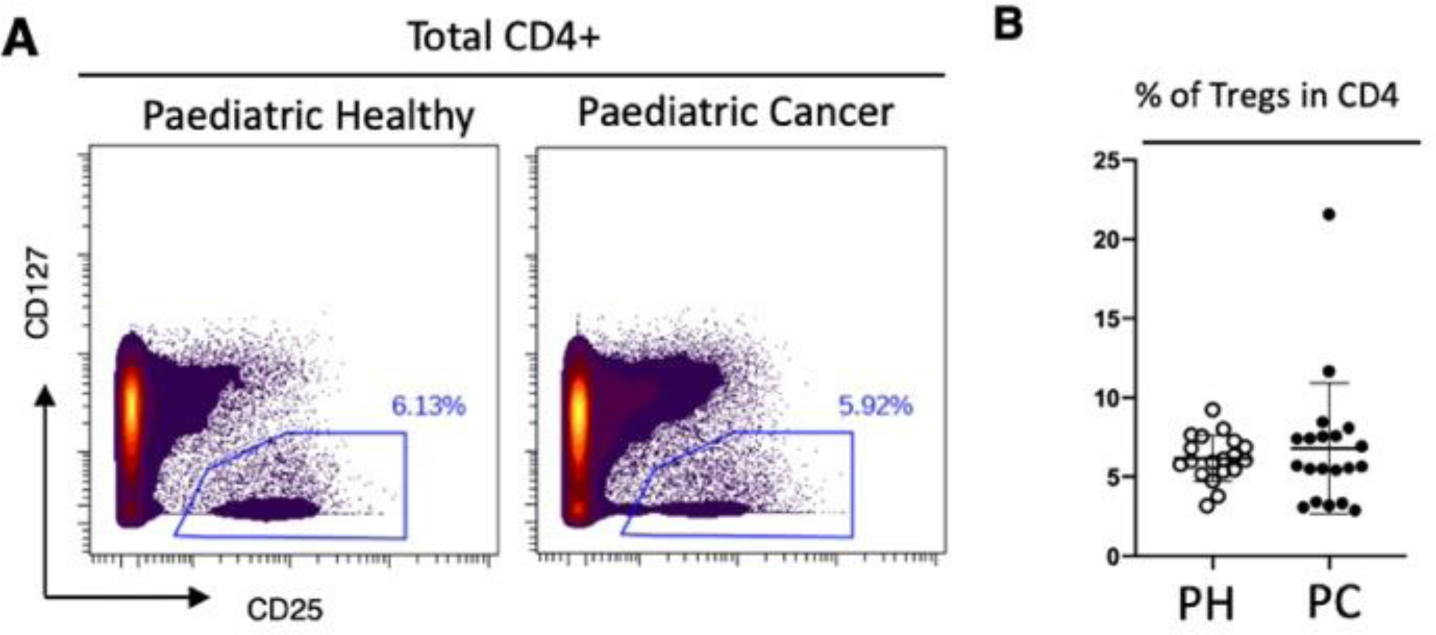
A) Gating strategy used to measure the frequency of CD25^hi^ CD127^lo^ regulatory T cells after gating on CD3^+^ CD4^+^ T cells from concantenated fcs files as show in SI figure 2. B) Frequency of CD25^hi^ CD127^lo^ regulatory T-cells for individuals. The error bars show the mean +/- one SD. Mann-Whitney test was used to compare the two groups with no significant difference.

**SI Fig 9.**
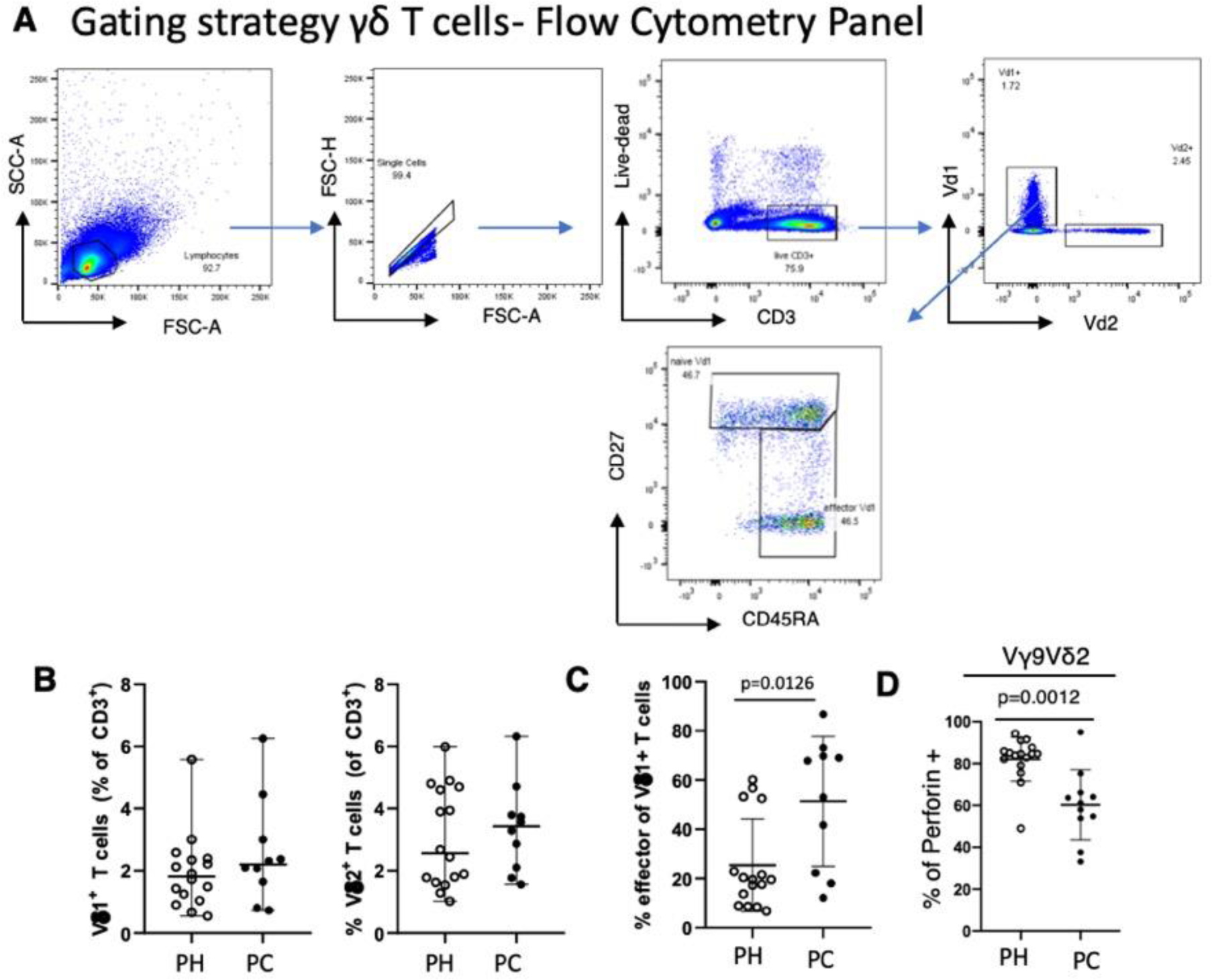
A) Gating strategy used to investigate gamma delta T cells and naïve and effector delta 1 (Vd1) T cells using a fluorescent flow cytometry antibody panel (SI Table 5). B) Frequencies of Vd1 and Vd2 (gamma delta 2) T-cells for individuals expressed as a percentage of CD3+ T cells. The error bars show the mean +/- one SD. The Mann-Whitney test was used to compare paediatric healthy (PH) and paediatric cancer (PC). C) Frequency of effector Vd1 T-cells (CD45RA+CD27-) is shown for individuals and is expressed as a percentage of total Vd1 T-cells. The error bars show the mean +/- one SD. Mann-Whitney test was used to compare PH and PC (p=0.0126). D) Frequency of perforin positive Vγ9Vδ2 T-cells is shown at the level of individuals and is expressed as a percentage of total Vγ9Vδ2. The error bars show the mean +/- one SD. Mann-Whitney test was used to compare PH and PC (p=0.0012).

**SI Fig 10.**
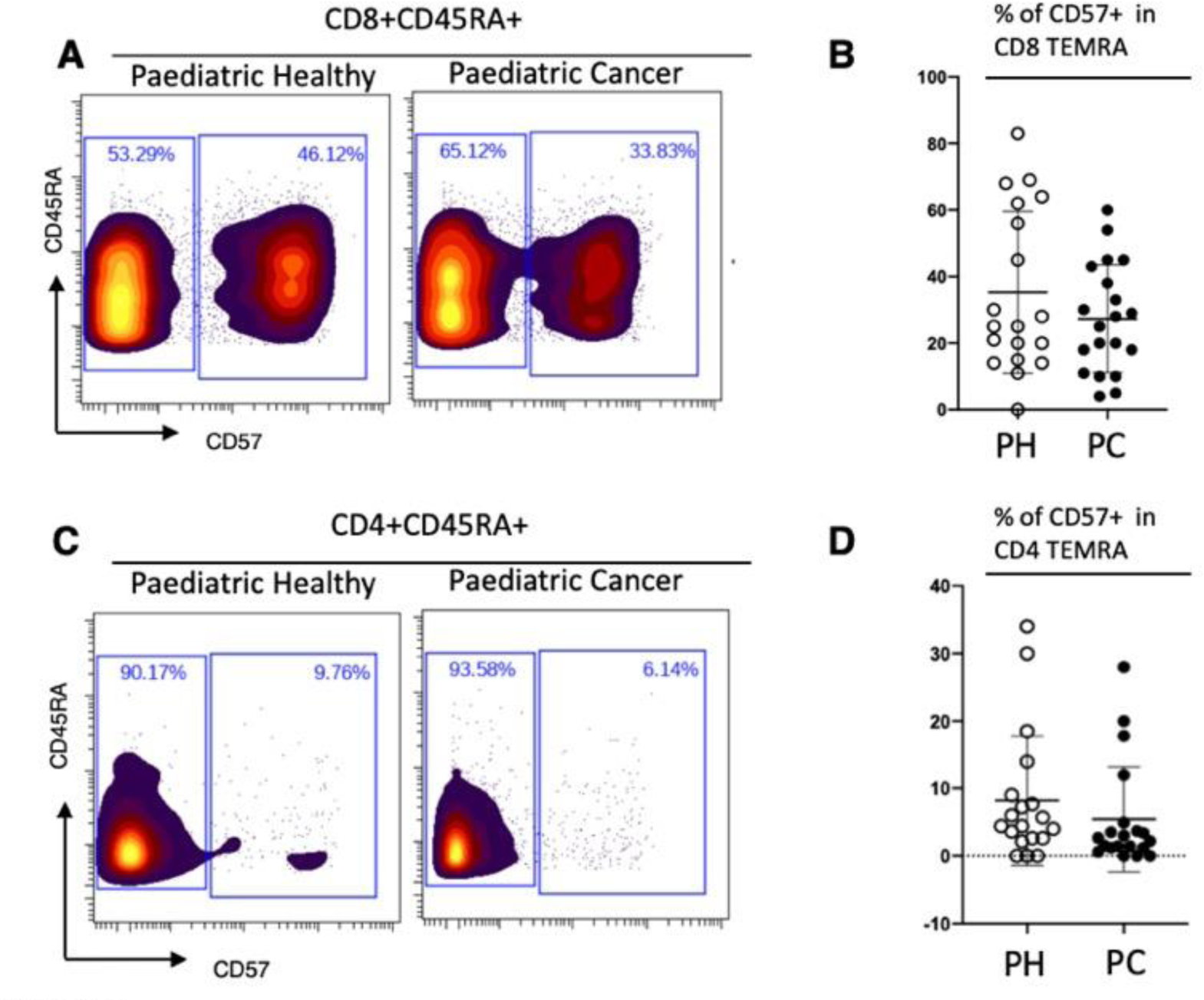
A) Gating strategy used to characterise CD57 expression on TEMRA T-cells using fcs files concatenated from healthy children or paediatric patients. B) Frequency of CD57+ CD8+ TEMRA cells for individual donors. The error bars show the mean +/- one SD. Mann-Whitney test was used to compare the two with no significant difference. C+D) The same analysis described for A and B applied to CD4+ T-cells.

## References

1. Board, I. of M. (US) and N. R. C. (US) N. C. P., Hewitt, M., Weiner, S. L. & Simone, J. V. The Epidemiology of Childhood Cancer. (National Academies Press (US), 2003).

2. Perkins, S. M., Shinohara, E. T., DeWees, T. & Frangoul, H. Outcome for Children with Metastatic Solid Tumors over the Last Four Decades. PLoS ONE 9, (2014).

3. Grabow, D. et al. The PanCareSurFup cohort of 83,333 five-year survivors of childhood cancer: a cohort from 12 European countries. Eur. J. Epidemiol. 33, 335–349 (2018).

4. Moreno-Vicente, J., Beers, S. A. & Gray, J. C. PD-1/PD-L1 blockade in paediatric cancers: What does the future hold? Cancer Lett. 457, 74–85 (2019).

5. Spitzer, M. H. et al. Systemic Immunity Is Required for Effective Cancer Immunotherapy. Cell 168, 487-502.e15 (2017).

6. Dowling, D. J. & Levy, O. Ontogeny of early life immunity. Trends Immunol. 35, 299–310 (2014).

7. Debock, I. & Flamand, V. Unbalanced Neonatal CD4(+) T-Cell Immunity. Front. Immunol. 5, 393 (2014).

8. Kohl, S. Human neonatal natural killer cell cytotoxicity function. Pediatr. Infect. Dis. J. 18, 635–637 (1999).

9. Olin, A. et al. Stereotypic Immune System Development in Newborn Children. Cell 174, 1277-1292.e14 (2018).

10. Zou, W. Regulatory T cells, tumour immunity and immunotherapy. Nat. Rev. Immunol. 6, 295–307 (2006).

11. Kagamu, H. et al. CD4+ T-cell Immunity in the Peripheral Blood Correlates with Response to Anti-PD-1 Therapy. Cancer Immunol. Res. 8, 334–344 (2019).

12. Emilie Mamessier, Lydie C. Pradel, Marie-Laure Thibult, Charlotte Drevet, Atika Zouine, Jocelyne Jacquemier, Gilles Houvenaeghel, François Bertucci, Daniel Birnbaum and Daniel Olive. Peripheral Blood NK Cells from Breast Cancer Patients Are Tumor-Induced Composite Subsets. J Immunol 1, 2424–36 (2013).

13. Wu, J. D. et al. Prevalent expression of the immunostimulatory MHC class I chain–related molecule is counteracted by shedding in prostate cancer. J. Clin. Invest. 114, 560–568 (2004).

14. Alexandrov, L. B. et al. Signatures of mutational processes in human cancer. Nature 500, 415–421 (2013).

15. Pfeiffer, M. et al. CD155 is involved in NK-cell mediated lysis of human hepatoblastoma in vitro. Front. Biosci. Elite Ed. 3, 1456–1466 (2011).

16. Raffaghello, L. et al. Expression and Functional Analysis of Human Leukocyte Antigen Class I Antigen-Processing Machinery in Medulloblastoma. Cancer Res. 67, 5471–5478 (2007).

17. Bernards, R., Dessain, S. K. & Weinberg, R. A. N-myc amplification causes down-modulation of MHC class I antigen expression in neuroblastoma. Cell 47, 667–674 (1986).

18. Borthwick, G. M., Hughes, L., Holmes, C. H., Davis, S. J. & Stirrat, G. M. Expression of class I and II major histocompatibility complex antigens in Wilms’ tumour and normal developing human kidney. Br. J. Cancer 58, 753–761 (1988).

19. Castriconi, R. et al. Natural Killer Cell-Mediated Killing of Freshly Isolated Neuroblastoma Cells: Critical Role of DNAX Accessory Molecule-1–Poliovirus Receptor Interaction. Cancer Res. 64, 9180–9184 (2004).

20. Ornatsky, O. et al. Highly multiparametric analysis by mass cytometry. J. Immunol. Methods 361, 1–20 (2010).

21. Laurens van de Maaten Geoffrey Hinton, G. H. Visualizing Data using t-SNE. J. Mach. Learn. Reasearch 9, 2579–2605 (2008).

22. McInnes, L., Healy, J., Saul, N. & Großberger, L. UMAP: Uniform Manifold Approximation and Projection. J. Open Source Softw. 3, 861 (2018).

23. Gassen, S. V. et al. FlowSOM: Using self-organizing maps for visualization and interpretation of cytometry data. Cytometry A 87, 636–645 (2015).

24. Diggins, K. E., Gandelman, J. S., Roe, C. E. & Irish, J. M. Generating Quantitative Cell Identity Labels with Marker Enrichment Modeling (MEM). Curr. Protoc. Cytom. 83, 10.21.1-10.21.28 (2018).

25. Narahara, K. et al. Regional mapping of catalase and Wilms tumor--aniridia, genitourinary abnormalities, and mental retardation triad loci to the chromosome segment 11p1305 p1306. Hum. Genet. 66, 181–185 (1984).

26. Mengos, A. E., Gastineau, D. A. & Gustafson, M. P. The CD14+HLA-DRlo/neg Monocyte: An Immunosuppressive Phenotype That Restrains Responses to Cancer Immunotherapy. Front. Immunol. 10, 1147 (2019).

27. Cooper, M. A., Fehniger, T. A. & Caligiuri, M. A. The biology of human natural killer-cell subsets. Trends Immunol. 22, 633–640 (2001).

28. Lee, S.-H., Fragoso, M. F. & Biron, C. A. A Novel Mechanism Bridging Innate and Adaptive Immunity: IL-12 Induction of CD25 to Form High Affinity IL-2 Receptors on NK Cells. J. Immunol. 189, 2712–2716 (2012).

29. Fu, B. et al. CD11b and CD27 reflect distinct population and functional specialization in human natural killer cells. Immunology 133, 350–359 (2011).

30. Erokhina, S. A. et al. HLA-DR+ NK cells are mostly characterized by less mature phenotype and high functional activity. Immunol. Cell Biol. 96, 212–228 (2018).

31. Scoville, S. D., Freud, A. G. & Caligiuri, M. A. Modeling Human Natural Killer Cell Development in the Era of Innate Lymphoid Cells. Front. Immunol. 8, (2017).

32. Ewelina Krzywinska. CD45 Isoform Profile Identifies Natural Killer (NK) Subsets with Differential Activity. PLoS ONE 11, e0150434 (2016).

33. Lopez-Vergès, S. et al. CD57 defines a functionally distinct population of mature NK cells in the human CD56dimCD16+ NK-cell subset. Blood 116, 3865–3874 (2010).

34. Somersalo, K., Tarkkanen, J., Patarroyo, M. & Saksela, E. Involvement of beta 2-integrins in the migration of human natural killer cells. J. Immunol. 149, 590–598 (1992).

35. Le Gars, M. et al. Pregnancy-Induced Alterations in NK Cell Phenotype and Function. Front. Immunol. 10, (2019).

36. Kurioka, A. et al. CD161 Defines a Functionally Distinct Subset of Pro-Inflammatory Natural Killer Cells. Front. Immunol. 9, 486 (2018).

37. Wendel, M., Galani, I. E., Suri-Payer, E. & Cerwenka, A. Natural Killer Cell Accumulation in Tumors Is Dependent on IFN-γ and CXCR3 Ligands. Cancer Res. 68, 8437 (2008).

38. Barrow, A. D. & Colonna, M. Exploiting NK Cell Surveillance Pathways for Cancer Therapy. Cancers 11, (2019).

39. Pegram, H. J., Andrews, D. M., Smyth, M. J., Darcy, P. K. & Kershaw, M. H. Activating and inhibitory receptors of natural killer cells. Immunol. Cell Biol. 89, 216–224 (2011).

40. Bassani, B. et al. Natural Killer Cells as Key Players of Tumor Progression and Angiogenesis: Old and Novel Tools to Divert Their Pro-Tumor Activities into Potent Anti-Tumor Effects. Cancers 11, (2019).

41. Nagel, J. E., Collins, G. D. & Adler, W. H. Spontaneous or natural killer cytotoxicity of K562 erythroleukemic cells in normal patients. Cancer Res. 41, 2284–2288 (1981).

42. Molfetta, R., Quatrini, L., Santoni, A. & Paolini, R. Regulation of NKG2D-Dependent NK Cell Functions: The Yin and the Yang of Receptor Endocytosis. Int. J. Mol. Sci. 18, (2017).

43. Sallusto, F., Lenig, D., Förster, R., Lipp, M. & Lanzavecchia, A. Two subsets of memory T lymphocytes with distinct homing potentials and effector functions. Nature 401, 708–712 (1999).

44. Davey, M. S. et al. The human Vd2+ T-cell compartment comprises distinct innate-like Vγ9+ and adaptive Vγ9-subsets. Nat. Commun. 9, (2018).

45. Mochizuki, K. et al. Various checkpoint molecules, and tumor-infiltrating lymphocytes in common pediatric solid tumors: Possibilities for novel immunotherapy. Pediatr. Hematol. Oncol. 36, 17–27 (2019).

46. Raffaghello, L. et al. Downregulation and/or release of NKG2D ligands as immune evasion strategy of human neuroblastoma. Neoplasia N. Y. N 6, 558–568 (2004).

47. Matusali, G. et al. Plasma levels of soluble MICA and ULBP2 are increased in children allergic to dust mites. J. Allergy Clin. Immunol. 130, 1003–1005 (2012).

48. Monaco, E. L. et al. Human Leukocyte Antigen E Contributes to Protect Tumor Cells from Lysis by Natural Killer Cells. Neoplasia 13, 822–830 (2011).

49. Bae, D. S., Hwang, Y. K. & Lee, J. K. Importance of NKG2D-NKG2D ligands interaction for cytolytic activity of natural killer cell. Cell. Immunol. 276, 122–127 (2012).

50. Xu, W. & Larbi, A. Markers of T Cell Senescence in Humans. Int. J. Mol. Sci. 18, (2017).

51. Yap, M. et al. Expansion of highly differentiated cytotoxic terminally differentiated effector memory CD8+ T cells in a subset of clinically stable kidney transplant recipients: a potential marker for late graft dysfunction. J. Am. Soc. Nephrol. 25, 1856–1868 (2014).

52. Pawelec, G. et al. Human immunosenescence: is it infectious? Immunol. Rev. 205, 257–268 (2005).

53. Kunert, A. et al. CD45RA+CCR7-CD8 T cells lacking co-stimulatory receptors demonstrate enhanced frequency in peripheral blood of NSCLC patients responding to nivolumab. J. Immunother. Cancer 7, 149 (2019).

54. Togashi, Y., Shitara, K. & Nishikawa, H. Regulatory T cells in cancer immunosuppression - implications for anticancer therapy. Nat. Rev. Clin. Oncol. 16, 356–371 (2019).

55. André, P. et al. Anti-NKG2A mAb Is a Checkpoint Inhibitor that Promotes Anti-tumor Immunity by Unleashing Both T and NK Cells. Cell 175, 1731-1743.e13 (2018).

56. Felices, M. et al. Continuous treatment with IL-15 exhausts human NK cells via a metabolic defect. JCI Insight 3, pii: 96219..

57. Wang, W., Erbe, A. K., Hank, J. A., Morris, Z. S. & Sondel, P. M. NK Cell-Mediated Antibody-Dependent Cellular Cytotoxicity in Cancer Immunotherapy. Front. Immunol. 6, (2015).

58. Federico, S. M. et al. A Pilot Trial of Humanized Anti-GD2 Monoclonal Antibody (hu14.18K322A) with Chemotherapy and Natural Killer Cells in Children with Recurrent/Refractory Neuroblastoma. Clin. Cancer Res. 23, 6441–6449 (2017).

59. Kreher, C. R., Dittrich, M. T., Guerkov, R., Boehm, B. O. & Tary-Lehmann, M. CD4+ and CD8+ cells in cryopreserved human PBMC maintain full functionality in cytokine ELISPOT assays. J. Immunol. Methods 278, 79–93 (2003).

